# Spatial Logic and Evolutionary Innovation in Human Placentation

**DOI:** 10.64898/2026.09.04.749511

**Authors:** Zhida Luo, Cheng Wang, Yan Zhou, Tuhin Kumar Guha, Purnima Narasimhan Iyer, Selina Lao Mason, Areca Smit, Xiaofei Sun, Michael P. Snyder, Gary M. Shaw, David K. Stevenson, Antonina Frolova, Sarah K. England, Virginia D. Winn, Susan J. Fisher, Jingjing Li

## Abstract

Human pregnancy unfolds in a unique mosaic tissue context. Embryonic/fetal placental and maternal uterine cells (decidua basalis) coalesce, forming the basal plate where immune tolerance is coordinated and the utero-placental circulation is initiated. The remaining approximately 70% of the maternal-fetal interface is comprised of the chorionic membranes, an epithelial-like layer of placental cells that lies adjacent to but does not coalesce with the overlying decidua capsularis and parietalis. It is unknown how these two regions of the maternal-fetal interface, which are comprised of seemingly similar cell types, diverge anatomically and functionally. Likewise, it is unknown whether the molecular characteristics of the maternal-fetal interface vary across human populations, potentially contributing to population disparities in pregnancy complications. Here, we generated large-scale paired single-nucleus RNA-seq and chromatin-accessibility profiles of the basal plate and chorionic membranes-associated decidual compartments from diverse-ancestry pregnancies, integrated with spatial transcriptomics and multiplexed protein imaging. We show that the two interfaces share a conserved cytotrophoblast differentiation hierarchy, which is deployed differently resulting in the observed distinct architectures. In the chorionic membranes, progenitor, early, and mature extravillous cytotrophoblast (EVT) states form an epithelial-like laminar shell. In the basal plate, this hierarchy is elaborated upon to enable deep placentation: EVTs that arise early in pregnancy invade the farthest into the uterus, while mature EVTs accumulate superficially to sculpt the local maternal immune microenvironment. Comparative analysis identified DSC4, a decidual stromal cell subtype, as an anti-invasive barrier that is densely enriched adjacent to the chorionic membranes but substantially reduced at the basal plate. Population-resolved analysis further identified the DSC4 marker *NID2*, which encodes the basement-membrane glycoprotein nidogen-2, as an ancestry-associated rheostat of EVT invasion. A specific *NID2* promoter haplotype arose on the modern-human lineage, is absent from available archaic-human genomes, and shows evidence of positive selection in Eurasian populations. This haplotype is associated with reduced local chromatin accessibility and lower *NID2* expression in DSC4 cells. Consistent with this genetic association, extracellular NID2 directly suppressed the invasion of primary human cytotrophoblasts *in vitro*. Together, these findings reveal how a shared developmental program is spatially reconfigured across distinct regions of the maternal–fetal interface and identify a recent modern-human regulatory innovation that modulates maternal decidual restraint of fetal trophoblast invasion across populations.

## Introduction

The human maternal-fetal interface is comprised of two distinct compartments. In one, the placenta invades the decidua basalis, forming the basal plate (BP). In the other, the chorionic membranes (CM) lie adjacent to, but do not invade, the decidua parietalis (Fig. 1A-C).^1–3^ These compartments have broadly similar cellular constituents, yet they execute sharply divergent anatomical and functional programs. In the BP (Fig. 1B), embryonic/fetal cytotrophoblasts of the placental origin differentiate into extravillous trophoblasts (EVTs) that deeply invade the decidua. In the process they engage stromal, immune, and vascular cells to anchor the placenta, promote immune tolerance and remodel spiral arteries into high-capacitance vessels that supply the utero-placental circulation, which sustains fetal growth.^2^ In contrast, the CM, which contains cytotrophoblast remnants of the placenta, forms a multi-layered epithelium that does not invade the decidua parietalis. (Fig. 1C).^1^ How the shared cellular framework of the BP and CM develops into distinct compartments remains poorly understood.

**Figure 1.**
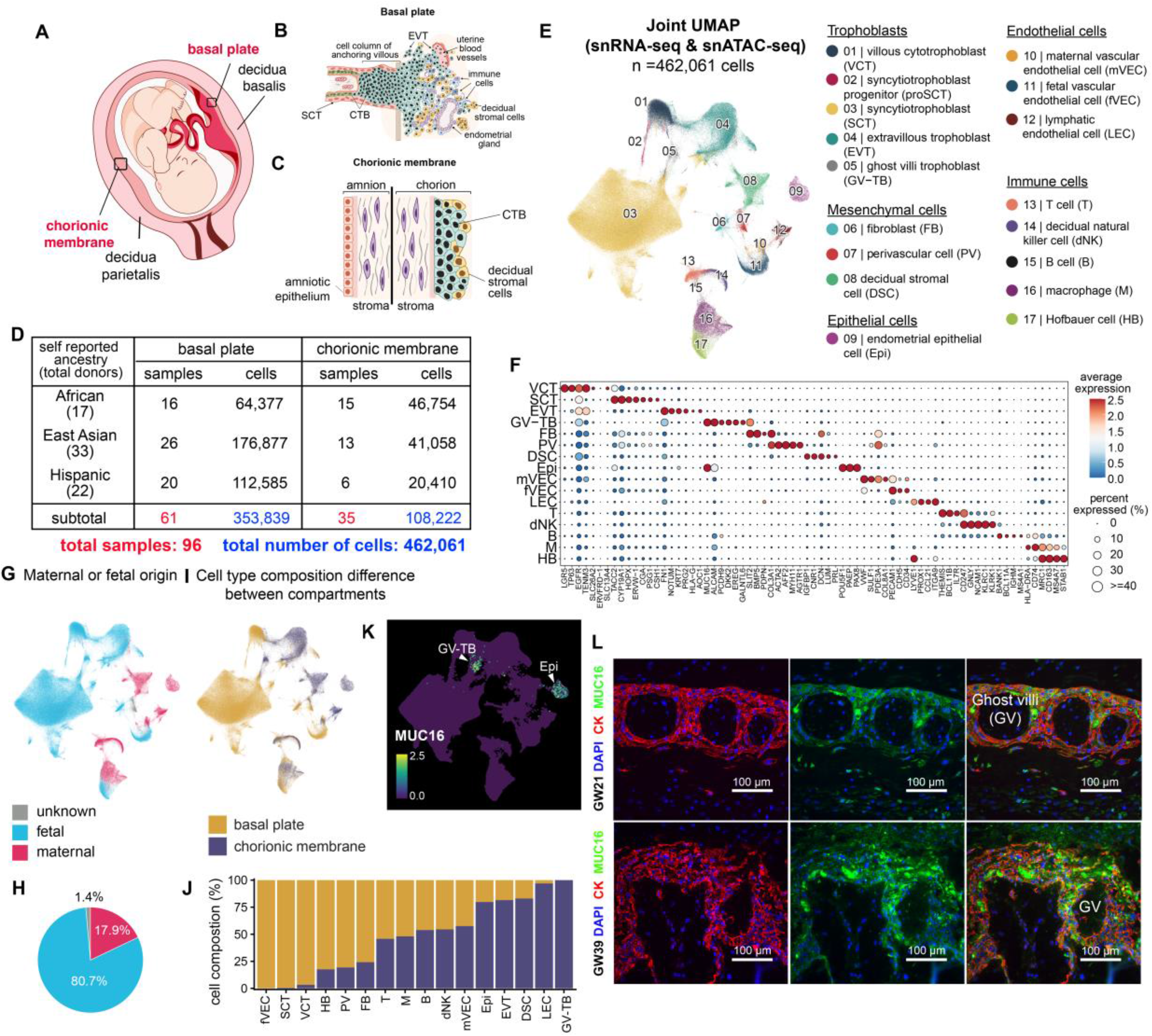
Single-nucleus multiome profiling of the basal plate and chorionic membranes. (A-C) Schematic overview of the pregnant uterus (A), basal plate (B) and chorionic membranes (C), showing their unique anatomy. SCT: syncytiotrophoblast; VCT: villous cytotrophoblasts; CTB: cytotrophoblast; EVT: extravillous cytotrophoblast. (D) Summary of the single-nucleus (sn) dataset by self-reported ancestry and tissue collection regions, showing the numbers of nuclei and samples from the basal plate and chorionic membrane samples. (E) Joint UMAP visualization of the sn-multiome data showing the major placental and uterine cell populations identified across the basal plate and chorionic membrane datasets. (F) Dot plot of representative marker genes used for cell-type annotation; color indicates average expression and point size indicates the percentage of cells expressing each gene. (G) UMAP visualization of fetal, maternal, and ambiguous cell origins. (H) Pie chart summarizing the overall proportions of fetal, maternal, and ambiguous cells. (I) UMAP visualization showing cells according to their tissue of origin. (J) Stacked bars showing the cell-type composition of the basal plate and chorionic membrane. (K) UMAP showing MUC16 expression, highlighting ghost-villus trophoblasts (GV-TB) and epithelial cells (Epi). (L) Immunofluorescence staining showing cytokeratin (CK; red), MUC16 (green), and DAPI nuclear staining (blue) in chorionic membranes at 21 and 39 gestational weeks. Merged images are shown in the right panels. Ghost villi (GV) are indicated.

Of the two, formation of the BP is best described. Here, EVT differentiation gives rise to extreme cellular heterogeneity (Fig. 1B). Our recent single-cell analysis of the BP revealed diverse EVT states distributed from the superficial to the deep decidua,^4^ suggesting that this diversity reflects functional specializations. The acquisition of EVT invasiveness has often been described as a partial epithelial-to-mesenchymal transition (EMT).^5,6^Yet several observations remain difficult to reconcile within this framework.^7^ From our recent single-cell data,^4^ invasive EVTs retain strong cytotrophoblast epithelial identity, including cytokeratin and E-cadherin expression (Fig. S1A), with limited acquisition of canonical mesenchymal markers such as Vimentin (Fig. S1A). Moreover, EMT-defining transcriptional factors, such as SNAI1/2, ZEB1/2, and TWIST1/2, are absent or only weakly expressed (Fig. S1A). At a global cell-state level, invasive EVTs occupy a transcriptomic space distinct from mesenchymal cells (Fig. S1A). These observations suggest that EVTs adopt a unique cytotrophoblast-specific invasive differentiation program. However, its mechanistic basis remains poorly understood.

Defective EVT invasion is one of the central mechanisms underlying major pregnancy complications.^8–11^ Insufficient invasion compromises spiral artery remodeling and placental perfusion, contributing to preeclampsia, ^10–13^ fetal growth restriction,^8,9^ recurrent pregnancy loss,^14^ and a subset of preterm births.^14^ Conversely, excessive or misdirected invasion underlies placenta accreta spectrum disorders, in which cytotrophoblasts penetrate too deeply into the uterine wall and fail to separate properly at birth, which can result in life-threatening bleeding.^15^ Most notably, these complications display striking population disparities. For example, preterm birth disproportionally affects African-ancestry populations, approximating 20% in Kenya^16^ and 14.7% among African Americans in the United States,^17^ in comparison to European- (∼10%)^17^ and East Asian-ancestry populations (∼5%).^18,19^ For decades, an explanation for population-based disparities in pregnancy outcomes has centered primarily on socioeconomic determinants.^20–22^ Our recent work also suggested potential genetic contributions.^23^ We reasoned if defective placental cytotrophoblast invasion is a common biological axis of pregnancy complications, then the BP and CP compartments can be used as comparators for identifying regulators of invasion that display population variation.

Here, we generated large-scale paired single-cell RNA-seq and ATAC-seq profiles of the human BP (Fig. 1B), CM (Fig. 1C) and the associated deciduas from individuals of diverse genetic ancestry. This integrated atlas identified previously unrecognized cytotrophoblast states in the CM, resolved EVT differentiation in the BP into spatially compartmentalized and functionally specialized states, and uncovered cellular programs underlying BP-CM functional divergence. These data suggest that cytotrophoblast differentiation in the CM stops short of giving rise to mature EVTs, resulting in an epithelial-like cellular arrangement. In contrast, invasion in the BP occurs in a step-wise fashion in which one EVT state locally remodels the decidua to prepare a permissive path for another state specialized for long-range penetration. Unexpectedly, the strongest population-associated cellular program was centered in a decidual subtype that suppresses EVT invasion, thereby tuning the permissiveness of the uterine microenvironment in the BP versus the CM. We show that this cell-type-specific differentiation is driven by recent positive evolutionary selection for a modern human-specific regulatory haplotype, absent in archaic humans, revealing an evolutionary innovation that reshaped the uterine control of EVT invasion. Overall, this study establishes a population-resolved single-cell atlas of the entire human maternal-fetal interface and reveals how recent human evolutionary history has shaped cell-state programs that may contribute to population differences in pregnancy outcomes.

## Results

We collected BP and CM samples (Fig. 1A-C) from individuals with normal pregnancies who delivered at term, using the same sample inclusion and exclusion criteria as described in our earlier work ^4^ (Methods). Samples with overt signs of infection, infarction, excessive calcifications, fetal chromosomal abnormalities, pregnancy complications, and donors with other notable pregnancy-associated or medical conditions were excluded (Table S1). These samples provide a reference for defining cell-state dynamics at the conclusion of normal pregnancy, while still capturing intermediate states along differentiation, maturation, and tissue-remodeling trajectories that remain active at term. Together with our previous extensive profiling of early, mid and term gestation BP samples,^4^ our focused profiling in this study of the entire maternal-fetal interface (e,g., matched BP and CM samples) at term enables reconstruction of a more complete cytotrophoblast differentiation landscape across human gestation. Moreover, the intentional inclusion of pregnancies from diverse ancestral backgrounds enables identification of population-associated variation in cell trajectories that may shape compartment-specific cytotrophoblast environments.

Each fresh-frozen sample was subjected to histological inspection before molecular profiling to confirm anatomical location and tissue integrity. Samples were validated by the presence of placental chorionic villi and extensive cytotrophoblast invasion in BPs, and preserved chorion-amnion architecture in CMs. Following quality control, 96 samples from individuals with diverse self-reported ancestries were included in downstream analyses (Fig. 1D).

### Single-cell profiling of the BP and CM compartments

Using the 10x Genomics platform, we generated paired single-nucleus transcriptomic (snRNA-seq) and chromatin-accessibility (snATAC-seq) profiles from the same nucleus in each fresh frozen sample, yielding high-quality data for the paired 462,061 single nuclei from 96 samples (Fig. 1D and Fig. S1B) after stringent quality control and batch effect removal (STAR Methods, Fig. S1C-F). Integrating snATAC-seq and snRNA-seq, the representation of individual nuclei in the joint UMAP space is shown in Fig. 1E, where cell types are unambiguously annotated by their respective markers as used in our previous study^4^ (Fig. 1F for gene expression and Fig. S1G for promoter accessibility of marker gene). These cell types were consistently represented across individuals from diverse ancestral backgrounds (Fig. S2A). Using our previous framework, cells were assigned as maternal or fetal in origin (Fig. 1G-H),^4^ with fetal identity independently confirmed by *UTY* expression in male-fetus pregnancies (Fig. S2B), Categorizing these cells by their anatomic origin from the BP or the CM enables identification of cell types preferentially located in one compartment relative to the other (Fig. 1I-J).

As expected, fetal endothelial cells showed strong compartment specificity for the BP, consistent with the vascularized villous architecture of the placenta. Conversely, lymphatic endothelial cells were strongly enriched in the CM compartment and depleted from the BP, consistent with prior observations.^24,25^ Although Hofbauer cells were most strongly enriched in the BP, consistent with the fact that these fetal macrophages reside in the placental villous stroma, a subset was also detected in the CM, which was confirmed by our spatial data (see below). This observation is consistent with prior reports of fetal-origin macrophages in the chorionic-amniotic membranes,^26,27^ but further shows that they share a close transcriptomic relationship with Hofbauer cells in the placenta. Within the fetal trophoblast lineage, villous cytotrophoblasts (VCTs) and syncytiotrophoblasts (SCTs) were strongly depleted in the CM but enriched in the BP, as expected from the presence of placental chorionic villi in the BP compartment (Fig. 1J). By contrast, EVT-like cytotrophoblasts were significantly enriched in the CM (Fig. 1J), suggesting that the multi-layered trophoblast shell of the CM predominantly adopts EVT-like transcriptional states (see below). Importantly, these compartment-enriched cell types were consistently observed across all ancestral populations (Fig. S2C). While most cell types and states were observed in our previous single cell datasets, we identified one novel trophoblast cell state (cell cluster 05, Fig. 1E), connecting the VCTs (cell cluster 01) to the SCTs (cell cluster 03, Fig. 1E), but transcriptomically divergent from the pre-fusion SCT progenitors (cell cluster 02, Fig. 1E). This cell state/subtype is a unique component of the CM vs. the BP (Fig. 1J). Gene expression analysis identified MUC16 as specifically marking this trophoblast cell state (in addition to epithelial cells, Fig. 1K). Immunolocalization of MUC16 and pan-cytokeratin (CK), which marks trophoblasts, shows that these novel cells reside in ghost villi of the CM (Fig. 1L). Given that these villi are thought to be chorionic villi that regressed early in development,^28^ we charted the development of these cells. Notably, ghost-villus trophoblasts did not map along the canonical VCT-to-SCT differentiation axis of floating villi, but instead localized near the branch point at which VCTs enter the EVT lineage (Fig. 1E). As EVT differentiation is initiated within cytotrophoblast cell columns of anchoring villi, this positioning raises the possibility that ghost villi are the remnants of these villi.

### Spatial localization of major cell types

With the major cell types defined, we next spatially mapped these populations in second-trimester CM and associated decidua samples from normal pregnancies and compared their organization with our recently generated BP spatial profiles from the same gestational window, generated on the same platform and by the same technician.^4^ The rationale for selecting second-trimester tissues was that the major cellular compartments are broadly shared with late gestation samples in the absence of the pronounced biomechanical and architectural changes associated with membrane maturation and labor onset. We first performed whole-slide single-cell multiplexed protein imaging with a post-QC 31-antibody CODEX panel^29,30^ (HuBMAP validated, Fig. 2A, Table S2) on a representative CM tissue section, followed by submicron-resolution spatial whole-transcriptome profiling on multiple CM sections. CODEX enabled cell-type spatial localization at the protein level, whereas spatial transcriptomics provided unbiased transcriptome-wide resolution to refine cellular states and identify spatially restricted molecular programs.

**Figure 2.**
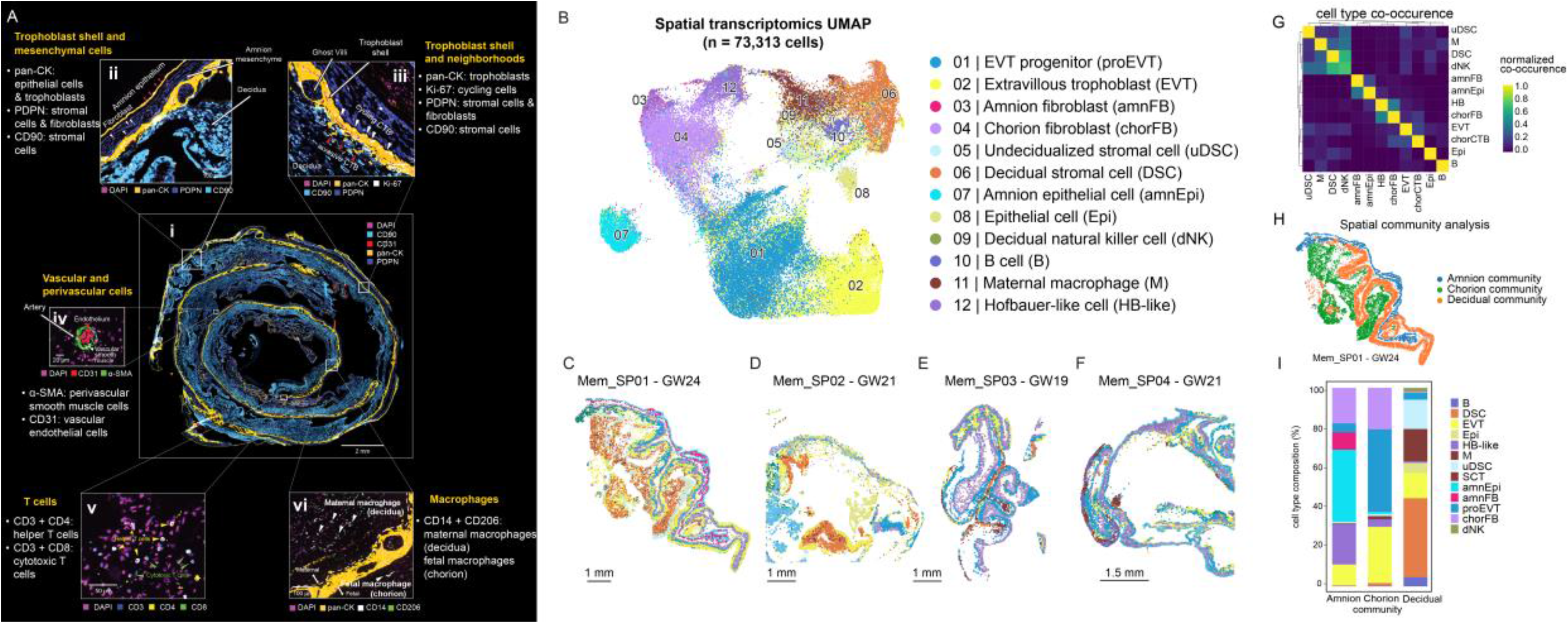
Spatial profiling of the human chorionic membrane. (A) CODEX imaging of a chorionic membrane (CM) sample comprised of a single layer of amniotic epithelial cells and the multi-layered cytotrophoblasts of the smooth chorion, separated by intervening fibroblasts and bounded at the periphery with the decidua. (i) Low-magnification overview of the CM. (ii, iii) Higher-magnification images of of the CM including ghost villi. (iv) Vascular/perivascular region highlighting endothelial and perivascular cells. (v) Immune cell localization showing T-cell subsets. (vi) Localization of maternal and fetal macrophages. (B) UMAP visualization identified by spatial transcriptomic profiling of chorionic membranes, showing 12 major cell types, including trophoblasts and fibroblasts as well as epithelial, vascular, and immune cell populations. (C-F) Spatial distribution of the major cell types across four chorionic membrane samples (GW19-24). (G) Clustered heatmap showing pairwise spatial co-occurrence among main cell types across all the four sections. (H) Spatial cell community analysis identified cell neighborhoods defining the amniotic, chorionic, and decidual compartments, exemplified by a GW24 sample. (I) Cell-type composition of these compartments as determined from the analysis of four chorionic membrane samples (GW 19-24).

Our CODEX panel was designed based on our prior experience with spatial profiling of the BP. We first prioritized antibodies that were validated and available through our HuBMAP CODEX workflow,^31,32^ and from this set selected markers that captured established cell types (Fig. 1E) as well as proteins of interest emerging from our transcriptomic analyses. Whole-slide imaging resolved the major anatomic compartments (Fig. 2A, i), which were then examined in detail. Notably, pan-CK-positive cytotrophoblasts formed a continuous shell at the CM surface. Discrete foci of EVT-like cells extended into the adjacent decidua where they were closely apposed to densely packed CD90-positive decidual cells (Fig. 2A, i-iii). They remained confined to localized niches rather than broadly infiltrating the decidua as in the BP. Thus, EVT invasion beyond the CM is spatially constrained. High-resolution imaging further identified cycling cytotrophoblasts (Ki-67-positive, Fig. 2A, iii) within the trophoblastic shell. In addition, CODEX imaging resolved organized maternal vascular (CD31^+^) and perivascular (α-SMA^+^) cells (Fig. 2A, iv), T-cell populations (CD3^+^/CD4^+^/CD8^+^; Fig. 2A, v), and macrophage-rich regions (CD14^+^ CD206^+^; Fig. 2A, vi), demonstrating that the CM contains highly structured epithelial, decidual, vascular, and immune microenvironments. Together, these data provide orthogonal protein-level validation of CM cellular architecture and identify focal sites where EVT-like cells emerge from the trophoblastic shell and invasion appears to be restricted by the decidua.

Having established the protein-level localization of major cell types by CODEX imaging, we next performed spatial transcriptomics to resolve the CM cell-state architecture at fine resolution. This approach allowed us to move beyond broad compartmental mapping and define how transcriptomically distinct cell states are spatially arranged across the CM-decidual interface, enabling direct comparison with our recently generated BP spatial atlas (on the same platform and by the same technician).^4^ Following our established spatial profiling workflow,^4^ we applied Stereo-seq^33^ to generate submicron-resolution whole-transcriptome maps of the CM. First, we immunostained the tissue sections to label cytotrophoblast (anti-pan-CK) and vascular (anti-CD31) compartments (Fig. S3A). After image-guided cell segmentation (Fig. S3B), spatially resolved transcript counts were aggregated within individual cell boundaries to generate cell-level whole-transcriptome profiles while preserving native tissue coordinates (Fig. S3C-E, QC details in Methods). Across four CM samples, we resolved the spatial transcriptomes of 73,313 cells (Fig. 2B–F) into major cell types based on the expression of biomarker genes (Fig. S3F), enabling their mapping within the native tissue architecture. The spatial distributions of these major cell types (Fig. 2C-F) showed strong concordance with the cell-type localization patterns independently defined by CODEX (Fig. 2A).

Leveraging these spatial transcriptomic data, we performed spatial neighborhood enrichment analysis to identify cell types that preferentially colocalize across the CM. In sharp contrast to the extensive heterotypic interactions observed in the BP,^4^ the CM exhibited a highly ordered architecture in which cells preferentially associated with others of the same type, indicating more limited cross-cell-type communication (Fig. 2G). Maternal decidual NK cells were a notable exception, showing extensive co-localization with both undecidualized and decidualized maternal stromal cells, as well as maternal macrophages, but minimal association with fetal cell populations (Fig. 2G).

We next performed spatial cell-community analysis to define recurrent “neighborhoods”, spatial niches composed of adjoining cells with characteristic cell-type compositions.^32,34^ This analysis identified three major communities corresponding to the principal anatomic compartments of the CM: the amnion, chorion, and decidua communities (Fig. 2H, Fig. S3G). Each community displayed a distinct cellular composition, exemplified by the selective enrichment of amniotic epithelial cells, cytotrophoblasts, and decidual stromal cells in the amnion, chorion, and decidual communities, respectively (Fig. 2I). Notably, although EVT-progenitor-like and EVTs constituted the dominant fetal trophoblast populations in the CM, only a small fraction of EVTs were assigned to the decidual community. These cells likely represent localized foci where EVTs have emerged from the CM but are spatially constrained from invading by the densely apposed decidual stroma (see below). Moving beyond broad cell-type characterizations, we next integrated the single-nucleus (Fig. 1E) and spatial datasets (Fig. 2B-F) to reconstruct the developmental trajectories and spatial organization of cytotrophoblasts at the entire maternal-fetal interface.

### Cytotrophoblast trajectories at the BP and CM interfaces

Focused analysis of BP and CM cytotrophoblasts resolved intermediate cell states connecting VCTs to compartment-specific trophoblast fates (Fig. 3A-D). In the BP (Fig. 3A), VCTs initiated an EVT differentiation program through progenitor EVT states marked by *ITGA2* and *KRT6A* (Fig. 3B). These cells subsequently progressed through early EVT states (*TCF7L2*-high, Fig. 3B) before maturing into late EVT populations characterized by strong *AOC1* expression (Fig. 3B, Fig. S4A), which marks mature EVTs.^4,35^ In addition to these canonical trophoblast states, profiling of term BP samples identified a previously unrecognized trophoblast population that predominantly appeared at term (Fig. 3A, cluster 05), and was rare or not identified in our prior first- and second-trimester BP datasets^4^ (Fig. S4B) as well as in previous single cell datasets.^36–38^ In our companion study, we show that this state is found to be selectively expanded in severe early-onset preeclampsia (PE). Because this population is characterized in detail in the companion study, it is not further discussed here.

**Figure 3.**
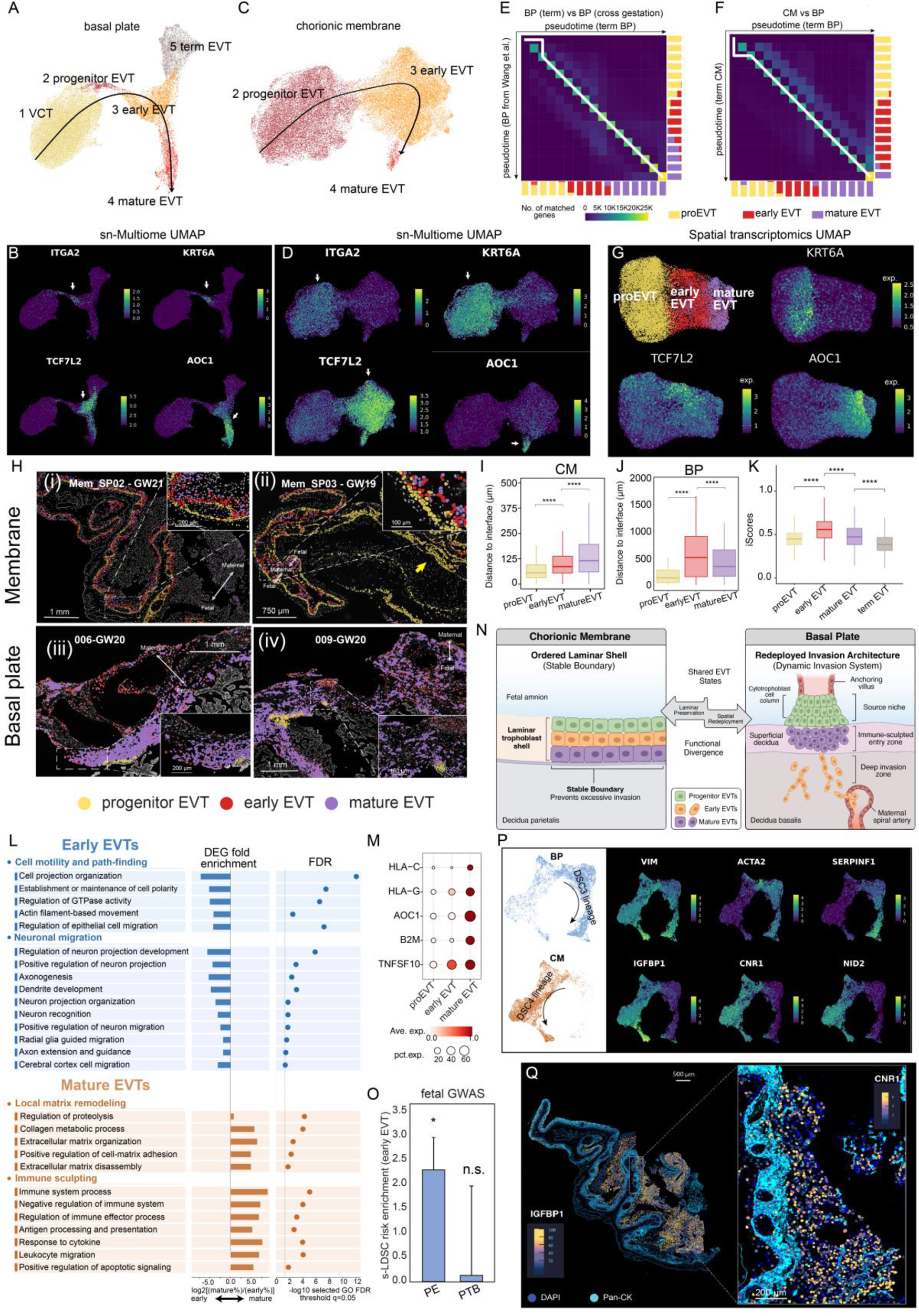
Distinct spatial organization and differentiation of cytotrophoblasts and decidual cells across the basal plate and chorionic membranes. (A-D) UMAPs and corresponding feature plots showing trophoblast subpopulations, inferred EVT differentiation trajectories, and selected marker expression in the basal plate (BP; A, B) and chorionic membranes (CM; C, D). (E-F) Alignment of pseudotime-ordered EVT populations across datasets. (E) BP cytotrophoblasts mapped to the published EVT developmental trajectory from Wang et al.^4^ (F) EVTs from the term CMs mapped to term BP EVTs. Heatmap color indicates the number of matched genes, and marginal annotations show the proportions of progenitor, early, and mature EVT cells. (G) UMAP visualization of CM cytotrophoblast and EVT populations identified from spatial transcriptomic data, with feature plots showing KRT6A, TCF7L2, and AOC1 expression across progenitor, early, and mature EVT states. (H) Spatial distribution of progenitor, early, and mature EVT populations across two CM samples (i) Mem_SP01 (GW 21) and (ii) Mem_SP03 (GW19) and two basal plate samples (iii) 006-GW20 and (iv) 009-GW20 from Wang et al.^4^ Enlarged regions provide detailed views of EVT subtype localization. Yellow arrows indicate regions where the trophoblast shell is not uniformly organized into three distinct layers, as described in the text. (I-J) Distances of progenitor, early, and mature EVT cells from tissue-specific reference boundaries: (I) the chorionic tissue boundary adjacent to the fetal chorionic fibroblast compartment in chorionic membrane samples, and (J) the trophoblast cell column in basal plate samples. (K) Boxplots show the distributions of invasiveness scores (iScores) across EVT subtypes in basal plate samples. (L) Gene Ontology biological process (GO-BP) enrichment analysis comparing BP early and mature EVT populations. Representative enriched terms are grouped by functional themes, and the directional DEG fold-enrichment plot indicates whether each process is preferentially associated with early EVT or mature EVT. (M) Expression of selected immune-related and EVT-associated genes across progenitor, early, and mature EVT populations. Color indicates scaled average expression, and point size represents the percentage of cells expressing each gene. (N) Schematic summary of divergent EVT organization across the CMs and BPs. Although cytotrophoblasts in both compartments share progenitor, early, and mature EVT differentiation states, they are organized in distinct tissue architectures. In the CM, EVTs are arranged as an ordered laminar trophoblast shell that forms a stable boundary and limits invasion. In the BP, EVTs are organized into successive zones that we theorize correspond to stages of invasion. Cytotrophoblast cell columns bridge the maternal–fetal interface, terminating in the superficial decidua, where they give rise to mature EVTs enriched in matrix-remodeling and immune functions. Less mature EVTs are enriched in pathfinding and migratory pathways that could facilitate deeper decidual invasion, where they remodel spiral arteries. (O) Bar plot showing stratified LD score regression (s-LDSC) enrichment of the early EVT gene set for preeclampsia (PE) and preterm birth (PTB) GWAS datasets. Error bars indicate standard errors; P < 0.05; n.s., not significant. (P) UMAP visualization of decidual lineages at the maternal-fetal interface showing that the CM is associated with the DSC4 lineage that comprises the decidua parietalis. The basal plate is dominated by the DSC3 lineage, which is a major component of the decidua basalis. Key DSC marker genes are indicated. (Q) Spatial distribution of decidual stromal cells in the fetal membrane sample Mem_SP01. Spatial transcriptomic expression of IGFBP1 delineates the broader DSC compartment, while CNR1 expression in the magnified region marks the DSC4 subtype. CNR1-expressing DSC4 cells are preferentially localized adjacent to the Pan-CK⁺ chorionic trophoblast.

The presence of the complete canonical EVT differentiation trajectory (Fig. 3A) suggests that the EVT differentiation continuum is well preserved in term BP samples. For further confirmation, we benchmarked this inferred VCT-to-EVT progression (arrows in Fig. 3A) to our recently published cross-gestational EVT differentiation trajectory^4^ using the pseudotime alignment algorithm (Methods).^39^ This alignment showed that near-perfect concordance between our term BP cytotrophoblast states and the reference developmental trajectory, with progenitor EVTs, *TCF7L2*-high early EVTs, and mature *AOC1*-high EVTs ordered sequentially along the same differentiation axis (Fig. 3E). This high trajectory concordance confirms that the profiled term samples faithfully captured the transcriptional dynamics of the full EVT differentiation continuum. We therefore projected each EVT state onto our previously generated second-trimester BP spatial transcriptomic atlas,^4^ allowing us to localize these developmentally anchored EVT states and define their spatial organization during active placentation and formation of the maternal-fetal interface, as discussed below.

In the CM, we identified three cell states, corresponding to the progenitor, early and late EVT populations in the BP, as indicated by their transcriptomic positions in the UMAP (Fig. 3C) space and by their respective markers (Fig. 3D). The transcriptomic correspondence between CM and BP cell states profiled in this study was further validated by our pseudotime alignment of cytotrophoblasts from both compartments (Fig. 3F). Despite the transcriptomic resemblance of each cell state, cell type composition was remarkably different between the BP and CM. In the BP, progenitor EVTs (marked by *ITGA2* or *KRT6A*, Fig. 3B) are rapidly depleted as they progress toward invasive EVT states. In sharp contrast, progenitor-like EVTs were substantially enriched in the CM (the same markers, Fig. 3D, Fig. S4C). This CM-specific expansion may reflect reduced differentiation efficiency and/or selective retention at this intermediate stage. Notably, as described below, our spatial mapping targeting the CM showed that these progenitor-like EVTs formed the basal layer of the chorionic trophoblast shell. Thus, the CM appears to leverage an expanded progenitor-like EVT population to build the CM while restricting full progression toward maturation.

To map CM trophoblast states *in situ*, we analyzed Stereo-seq spatial transcriptomes of fetal cytotrophoblasts within the CK-immunostained trophoblast shell (Fig. 2C-F), excluding CK-positive epithelial cells from endometrial glands (Fig. S3A). Focusing on cytotrophoblasts, we mapped their single-cell transcriptomes onto the UMAP space, resolving three CM trophoblast states that matched the three EVT states defined in our single-nucleus atlas (Fig. 3A): *ITGA2/KRT6A*-positive progenitor EVTs, *TCF7L2*-high early EVTs, and *AOC1*-high mature EVTs (Fig. 3G). Projecting these cell states from the UMAP space back to their native tissue context revealed that these states were organized in a laminar sequence across the CM trophoblast shell (Fig. 3H, upper panels), with progenitor EVTs forming the basal layer facing the amnion (yellow), early EVTs positioned in the middle (red), and mature EVTs (purple) occupying the outermost maternal-facing layer adjacent to the decidua. Aggregating cells across all samples, we next quantified each one’s physical distance from the underlying chorionic fibroblasts (Fig. S4D). Consistent with the observed laminar organization (Fig. 3H, upper panels), physical distance from this boundary increased progressively with EVT maturation: progenitor EVTs were most proximal, early EVTs were positioned at intermediate distances, and mature EVTs were located nearest to the CM-decidual interface (Fig. 3I).

Notably, the epithelial-like CM trophoblast shell is not a uniformly organized three-layered structure. While the fetal-facing progenitor-like trophoblast layer is continuous, the early EVT-like and mature EVT-like layers are regionally discontinuous and locally absent in some areas (Fig. 3H, ii, yellow arrow). This pattern may reflect regional differences where EVT differentiation is incomplete, raising the possibility that these areas could be more vulnerable to rupture. Taken together, these findings at single cell resolution extend the conventional view of the CM trophoblast shell as a homogeneous chorionic trophoblast barrier by revealing a previously unrecognized laminar differentiation architecture, suggesting that the CM uses compartmentalized trophoblast maturation to establish a stable fetal-maternal boundary.

### Spatial redeployment of EVT states in the BP

Progenitor, early, and mature EVT states in term BP showed strong transcriptomic correspondence with their respective counterparts in the published cross-gestational reference dataset (Fig. 3E) and were also consistently identified across CM and term BP samples (Fig. 3F). Therefore, we used these aligned transcriptomic states as developmental anchors and projected them onto matched second-trimester spatial maps of the CM generated in this study (Fig. 2B) and the BP profiled in our prior study.^4^ This strategy allowed us to leverage the 2^nd^ trimester samples, when EVT invasion and decidual remodeling are active in the BP, to ask whether the same EVT states occupy analogous anatomical positions in both compartments.

In the BP, progenitor EVTs were located adjacent to cell columns (yellow, Fig. 3H, lower panels), consistent with our previous observations.^4^ In contrast to the CM, which preserved a shell-like laminar organization of progenitor, early, and mature EVT states, the BP broke this spatial order and redeployed EVT states across the decidua. Mature EVTs (purple) no longer formed an orderly shell but invaded the superficial decidua. In contrast, early EVTs migrated further and reached the deepest decidual regions (red, Fig. 3H, lower panels). Quantitative spatial analysis further supported this compartment-specific organization. In the CM, EVT maturation was associated with increasing distance from the manually defined chorionic tissue boundary adjacent to the fetal chorionic fibroblast compartment (Fig. 3I), consistent with an ordered progression from progenitor-like cytotrophoblasts toward more differentiated EVT-like states. In contrast, analysis of all 16 BP samples from our spatial atlas^4^ revealed that this spatial organization was reversed. Early EVTs occupied the deepest portions of the decidua, with a median distance of approximately 500 μm from the maternal-fetal interface, whereas mature EVTs were positioned more superficially, at a median distance of approximately 300 μm (Fig. 3J; adjusted P < 1.49 x10^−147^, Wilcoxon rank-sum test). This spatial organization is consistent with our previous single-cell analysis, which showed that early-arising EVTs along the developmental trajectory exhibit the strongest invasiveness^4^. For additional confirmation, we computed the EVT invasiveness scores^4^ (iScores, see Method) for each EVT cell in this independently generated dataset. Indeed, the early EVTs had the highest iScores among all EVT populations (Fig. 3K). Statistical significance remained after stratifying the cells by sample, indicating that this result was not skewed by person-to-person variations (Fig. S4E). Thus, the orderly epithelial-like arrangement of differentiating EVTs in the CM is upended in the BP where early EVT states penetrate farthest into the decidua, while mature EVTs occupy the superficial decidua (Fig. 3N).

To define the molecular basis of the reversed positioning of early and mature EVTs in the BP, we focused on differentially expressed genes between the two populations (Methods). Genes significantly up-regulated in early EVTs (Table S3) were enriched for GO-BP terms related to cell morphogenesis, projection organization, actin cytoskeleton remodeling, small-GTPase signaling, cell polarity, epithelial cell migration, and developmental pathfinding (Fig. 3L, and Table S4), indicating a differentiation program aimed at the acquisition of directional motility, a cell migration mechanism deployed by (among other cell types) neurons. Indeed, the enriched terms significantly involved key neuronal migration processes, including positive regulation of neuron migration (FDR=0.018) and axon development (FDR=8.62 x10^−4^, Fig. 3L). In accord with this observation, many early EVT-upregulated genes are well-established regulators of neuronal migration, axon guidance, or developmental pathfinding, including *DSCAM*, *RND3*, *SEMA5A*, *SEMA6D*, *EFNA5*, *TENM3*, *DOCK3*, and *NEDD9* (Table S3). This points to the use of conserved guidance machinery to achieve deep decidual invasion, linking extracellular positional cues to polarity, protrusion formation, adhesion dynamics, and cytoskeletal remodeling. These data suggest that early-stage EVTs achieve deep decidual invasion by means of a sophisticated navigation system, in which decidual microenvironmental cues direct long-range interstitial migration toward deep tissue compartments. Given that ectopic pregnancy can result in live birth, it is likely that this navigation system can, at some level, operate autonomously.

In contrast, genes up-regulated in mature EVTs (Table S3), were enriched for cell-matrix adhesion, proteolysis, collagen metabolism, and wound-healing programs, indicating a shift from long-range directional migration toward local tissue remodeling (Fig. 3L and Table S4) consistent with the preferential accumulation of mature EVTs in the superficial decidua (Fig. 3H, panel iii and iv, cells purple). These cells also acquired a strong immune signature, marked by significant enrichment for cytokine response, antigen processing and presentation, leukocyte-associated pathways, and apoptotic/cell-death signaling (Fig. 3L). This immune transition was exemplified by pronounced upregulation of *HLA-C*, *HLA-G*, *AOC1*, *B2M*, and *TNFSF10*/TRAIL in mature EVTs (Fig. 3M). Because maternal immune cells broadly expressed the TRAIL death receptors TNFRSF10A/DR4 and TNFRSF10B/DR5 (Fig. S4F), these data support a model in which mature EVTs sculpt the superficial decidual niche by coupling HLA-associated immune modulation and evasion with TRAIL-dependent immune-cell elimination. Consistent with this model, our previous spatial analysis revealed pronounced compartmentalization of the decidua, with the superficial compartment markedly depleted of maternal immune cells, in sharp contrast to the immune-rich deep decidua.^4^ Together, these findings suggest that EVTs, which arise early in the differentiation trajectory, are specialized for deep uterine invasion. The mature EVT population, which arises later, may follow as far as the superficial BP where these cells stabilize this region of the maternal-fetal interface via matrix remodeling and immune regulatory programs (Fig. 3N).

Having defined distinct early and mature EVT states, we next investigated their clinical relevance by focusing on preeclampsia, a pregnancy complication characterized by impaired EVT invasion. Using summary statistics from a large-scale genome-wide association study of preeclampsia based on more than 6000 fetal genomes,^40^ we performed stratified linkage disequilibrium score regression (S-LDSC, Methods) using gene sets upregulated in early and mature EVTs, respectively (Table S5). Fetal genetic risk for preeclampsia was significantly enriched in genes preferentially expressed by the deep-penetrating early EVT population (Fig. 3O), but not in genes associated with the more superficial mature EVT state (Table S5). The results were robust to SNP inclusion windows, remaining unchanged when considering either gene-body SNPs or a ±10 kb flanking window (Table S5). As a control experiments, we applied the same analysis to an independent fetal-genome GWAS datasets for preterm birth before 37 weeks of gestation.^41^ The PTB dataset showed did not show any significance in either early or mature EVT genes (Fig. 3O; Table S5), indicating that the observed association is specific to fetal genetic risk for preeclampsia. Together, these findings identify early EVTs as the principal EVT population through which fetal genetic susceptibility to preeclampsia is manifested.

Lastly, despite the overall transcriptomic similarity of corresponding early and mature EVT states between the BP and CM, we asked whether individual genes marking these states exhibited compartment-specific differential expression, potentially reflecting functional specializations imposed by their distinct tissue environments. In early EVTs of the CM, we observed the strongest up-regulation of genes enriched for hypoxia responses (Table S6 and Table S7), which was consistent with its lack of a maternal blood supply. In mature EVTs, we observed the strongest up-regulation of genes enriched in cell junctions, suggesting their role in formation of a barrier (Table S7). Most notably, TRAIL expression was further elevated in mature EVTs of the CM relative to their BP counterparts (Table S6), suggesting enhanced suppression of maternal immune-cell infiltration. Thus, these comparative molecular features indicate that mature EVTs in the CM adopt enhanced barrier and immune suppression programs that inhibit interactions with the decidua parietalis. This is contrast to EVTs in the BP, which cooperate to carry out collective migration of the decidua basalis, which entails navigating deep decidual invasion, and superficially, wound healing and immune functions.

### Decidual architectures diverges between the BP and CM

EVTs of the CM were tightly encased by maternal decidual cells (Fig. 2A, iii), suggesting that invasion is actively constrained. Therefore, we investigated the possibility that the decidua parietalis forms a barrier to EVT invasion. Our recent BP analysis^4^ identified two major decidual cell (DSC) lineages arising from stromal cells: a DSC3 state enriched in the deep decidua and around uterine vessels, and a DSC4 state that localized to the superficial decidua.^4^ We also showed that decidualized DSC4 cells suppress EVT invasion through endocannabinoid signaling.^4^ Here, we mapped decidual cells from the BP and CM samples in this study to the DSC3 and DSC4 lineages identified in our previous work^4^ Specifically, we used *VIM* expression to confirm their stromal identity, *ACTA2* to mark undecidualized cells and *SERPINEF1* positivity to identify the DSC3 lineage. The DSC4 lineage co-expressed *IGFBP1*, *CNR1* and *NID2* (Fig. 3P, right panel). Comparing snRNA-seq data regarding the decidual cell composition of the basalis vs. parietalis unexpectedly revealed that decidual cells in a DSC4-like state adjoined the CM (Fig. 3P). This finding was independently confirmed by spatial transcriptomics (Fig. 3Q and Fig. S4G), which revealed a continuous *IGFBP1*+*CNR1*+ DSC4-rich layer surrounding the EVT shell. Given the established anti-invasive function of DSC4 cells,^4^ they may form a barrier that confines EVT invasion. This is in contrast to DSC4 cells in the BP that are also found in the superficial decidua but are dismantled into a discontinuous sparse layer frequently positioned adjacent to the termini of cytotrophoblast cell columns^4^. In the context of the CM results, the cellular architecture in the BP may create routes for EVTs to invade the decidua.

### Positive selection shapes population differences in DSC identity

Given the potential contribution of population genetic variation to placental physiology, we next examined whether genetic ancestry was associated with cell identity or functional state. Rather than relying solely on self-reported ancestry, we inferred genetic ancestry directly from sequencing data. For each sample, we first assigned single-cell reads to maternal or fetal origin (Fig. 1G), enabling separate reconstruction of maternal and fetal genetic variation. We then performed single-cell SNV calling using Monopogen,^42^ a method specifically designed and benchmarked for variant detection from single-cell sequencing data. This allowed us to derive resolvable genotypes for both maternal and fetal genomes in each pregnancy. By aggregating genotype information across individuals, we estimated ancestry proportions relative to African, European, and East Asian reference populations from the 1000 Genomes Project (Methods). We focused on these three reference populations because they captured the major axes of genetic variation represented in our cohort (Fig. S5A). Importantly, these references were not used to assign individuals to discrete racial or ancestry categories, but rather as population anchors for estimating continuous ancestry coefficients (Fig. 4A; Methods), allowing ancestry-associated cellular variation to be modeled quantitatively. We next assessed how well genetically inferred ancestry corresponded with maternal self-reported ancestry available in our cohort. Overall, the two measures showed broad concordance (Fig. 4A), supporting the validity of the genotype-based inference. For the small subset of discordant cases, we used genetically inferred ancestry in downstream analyses, because it provides a quantitative and directly measured estimate of ancestry from the sequencing data.

**Figure 4.**
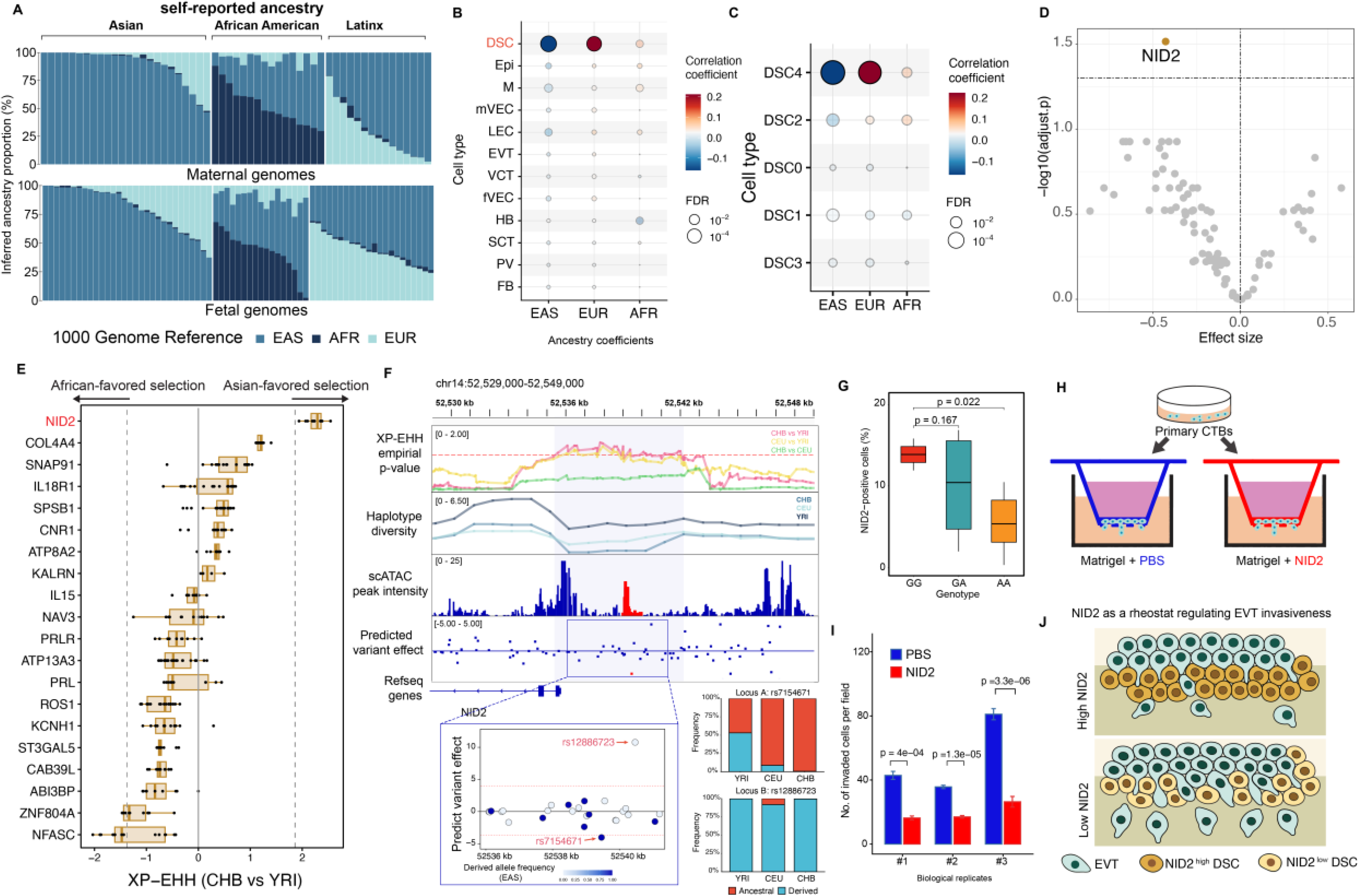
Genetic ancestry-associated regulation of *NID2* and its impact on trophoblast invasion. (A) Genetic ancestry composition of fetal and maternal genomes. Stacked bars show inferred EAS, AFR, and EUR ancestry proportions for individuals grouped by self-identified Asian, African American, or Latinx identity. (B) Multivariable linear regression of cell-type identity scores on EAS, EUR, and AFR ancestry proportions using Equation 1, adjusting for clinical and technical covariates. The x-axis indicates the corresponding ancestry regression coefficient, and point size represents the FDR-adjusted P value. (C) Multivariable linear regression of DSC subtype identity scores using Equation 1, adjusting for clinical and technical covariates. The x-axis indicates the corresponding ancestry regression coefficient, and point size represents the FDR-adjusted P value. (D) Multivariable linear regression of DSC marker-gene expression using Equation 2, adjusting for clinical and technical covariates as well as compositional dependence among ancestry components. Effect sizes are plotted against FDR-adjusted significance. Dashed lines indicate an effect size of zero and the significance threshold. (E) ХР-ЕНН scores (CHB versus YRI) for SNPs within the promoter regions of DSC4 marker genes. Positive values indicate Asian-favored selection, whereas negative values indicate African-favored selection. Dashed lines indicate the empirical top 5% thresholds of the XP-EHH score distribution. (F) Genome-browser view of the *NID2* promoter region showing XP-EHH empirical P values, Shannon entropy, snATAC-seq accessibility, predicted variant effects, and RefSeq gene annotations. The shaded area marks the selection-associated interval. The enlarged panel highlights two variants within this region that fall within the top 5% of predicted regulatory effects. rs7154671 is predicted to reduce chromatin accessibility, whereas rs12886723 is predicted to increase chromatin accessibility. The right panels show the ancestral and derived allele frequencies of both variants in YRI, CEU, and CHB populations. (G) Percentage of *NID2*-positive cells across inferred rs7154671 genotype groups in the single-nuclei dataset. (H) Schematic of the experimental design for functional validation of NID2 using primary cytotrophoblasts. Cells were treated with PBS or recombinant NID2 and subjected to Matrigel invasion assays. (I) Quantification of cytotrophoblast invasion following PBS or recombinant NID2 treatment. Invaded cells were quantified across three independent experiments. (J) Proposed model summarizing the relationship between NID2 expression in decidual stromal cells (DSCs) and extravillous trophoblast (EVT) invasion. High NID2 expression in DSCs creates a more restrictive decidual environment and limits EVT invasion, whereas lower NID2 expression reduces this barrier and permits deeper or more extensive EVT invasion.

We sought to determine whether cell identity or functional state varies with genetic ancestry. For each cell type or states, we first calculated cell identity/state signature for individual cells using top 100 highly specific marker genes (Fig. S5B). These scores were then summarized at the donor level by taking the median across all corresponding cells from each individual, yielding one score per donor per cell type or state. This aggregation treated the individual, rather than the single cell, as the statistical unit and reduced the impact of sparsity in rare, heterogeneous, or transitional populations. Donor-level scores were then correlated with continuous ancestry coefficients, with each ancestry component tested separately. Available biological and technical covariates were incorporated into the association models to assess whether ancestry-associated cellular variation remained after accounting for potential confounding factors (Methods). Because fetal and maternal cells originate from distinct genomes, fetal ancestry coefficients were used for trophoblasts, Hofbauer cells, and other fetal populations, whereas maternal ancestry coefficients were used for decidual stromal, maternal immune cells and other maternal-derived populations. This analysis served as a screening strategy to nominate ancestry-associated cellular programs for subsequent population genetic analyses and functional validation.

Among all the broadly defined cell types identified from BP or CM, decidual stromal cells (DSCs) was the only cell type displaying significant association between cell type identity and population ancestries. Specifically, DSC identity scores displayed a significant negative association with East Asian ancestry after adjusting donor-level clinical and technical covariates (FDR=7.16e-06; Fig. 4B; Table S8a; Equation 1, Methods), but a significant positive association with European ancestry (FDR=4.33 × 10^−5^; Fig. 4B; Table S8a). In contrast, the association with African ancestry was close to zero (Table S8a). To account for the compositional dependence among the three ancestry components, we further regressed DSC identity scores on East Asian ancestry, the within-individual difference between European and African ancestry, and donor-level covariates (Equation 2, Methods). This parameterization tested whether the contrast between the European and African ancestry components was associated with cell identity scores independently of the East Asian component. Again, the negative association between East Asian ancestry and DSC identity remained significant (FDR=2.45 x 10^−6^; Table S8b), whereas the European-African contrast was not significant (FDR=0.589). These results indicate that the ancestry-associated variation in DSC identity is statistically driven by the East Asian ancestry component, with no detectable difference between European and African ancestry after adjustment. We therefore prioritized decidual stromal cells for downstream analyses of the genetic basis and molecular programs underlying ancestry-associated variation in DSC identity/state.

We next asked whether the ancestry association observed for the broad DSC compartment reflected a general shift across all decidual stromal states or was localized to a specific DSC subtype. We therefore calculated identity scores for each of the five DSC subtypes, DSC0 through DSC4 (Fig. 3P), using the same framework applied to the broad cell-type identity scores and repeated the covariate-adjusted ancestry association analyses described above (Methods). Strikingly, DSC4 recapitulated the ancestry-association pattern observed for the overall DSC compartment. In the single-component models (Equation 1, Methods), DSC4 identity scores were significantly negatively associated with East Asian ancestry and positively associated with European ancestry, whereas no significant association was detected with African ancestry (Fig. 4C; Table S8c). After accounting for the compositional dependence among ancestry components, the negative association between East Asian ancestry and DSC4 identity remained significant, whereas the European-minus-African ancestry contrast was not significant (Table S8d). None of the other DSC subtypes showed a significant association with ancestry after correction for multiple testing. These results indicate that the ancestry-associated variation observed for the broad DSC compartment is predominantly attributable to the DSC4 state and is best characterized as an East Asian-versus-non-East-Asian contrast, rather than a difference between European and African ancestry. Thus, genetic ancestry is not associated with a uniform alteration of the decidual stromal lineage but with selective variation in the representation or molecular identity of the superficial DSC4 subtype.

To identify the molecular program underlying the ancestry-associated variation in DSC4 identity, we extended the same donor-level association framework to the top 100 genes defining the DSC4 state (Methods). For each marker gene, we tested whether its expression across donors was associated with individual ancestry components using the single-component models in Equation 1 (Methods) and then evaluated the robustness of the association using the composition-aware parameterization in Equation 2 (Methods). Among the DSC4 marker genes, *NID2* showed the strongest negative association with East Asian ancestry and was the only marker that remained significant after correction for multiple testing (Fig. 4D; Table S8e–g). This association persisted after accounting for the compositional dependence among ancestry components: *NID2* expression remained significantly associated with the East Asian-versus-average-European/African ancestry contrast, whereas the European-minus-African contrast was not significant (Equation 2, Methods; Table S8h). Thus, the ancestry-associated reduction in DSC4 identity is accompanied by a concordant reduction in *NID2* expression and is again best characterized as an East Asian-versus-non-East-Asian pattern. These findings nominate NID2 as a principal molecular feature of the ancestry-associated DSC4 state and motivated subsequent investigation of the genetic variation regulating *NID2* expression.

Population differentiation can arise through neutral demographic processes without clear biological consequences, or through positive selection that drives local adaptation during human evolution and migration.^43,44^ We next tested whether the observed population ancestry effects were driven by evolutionary positive selection due to local adaptation during human population differentiation. For this analysis, we used the 1000 Genomes CHB population (Han Chinese in Beijing) as a representative East Asian reference population with well-characterized phased haplotypes and compared it with CEU (Utah residents with Northern and Western European ancestry) and YRI (Yoruba in Ibadan, Nigeria), the latter representing an African population with relatively limited recent non-African admixture, using XP-EHH (Methods).^45^ XP-EHH detects recent population-specific positive selection by detecting unusually extended haplotype homozygosity around an allele in one population relative to another, consistent with rapid allele-frequency increase before recombination has eroded the surrounding haplotype.^45^ This haplotype-based scan identified a significant selection signal at *NID2* (nidogen-2), a core marker of DSC4 (Fig. 3P), ranking within the top 5% of genome-wide XP-EHH scores and showing markedly extended haplotype homozygosity across the NID2 locus in East Asians relative to Africans (Fig. 4E). We also examined CHB-CEU and CEU-YRI comparisons to characterize how the *NID2* selection signal varied across population contrast (Fig. S5C-D). Overall, the signal was strongest in CHB vs YRI, remained evident in CEU vs YRI, and showed little differentiation between CHB and CEU.

### Fine-mapping of the selected functional variant at the NID2 promoter

Fine-scale examination of the significant XP-EHH signal localized the putative target of selection to the promoter-proximal region of *NID2*, displaying strong chromatin accessibility in DSC4 cells (Fig. 4F), where XP-EHH peak signals were observed in both East Asian (CHB) vs African (YRI) and European (CEU) vs African (YRI) comparisons (Fig. 4F, panel 1). The most parsimonious interpretation therefore supports positive selection specific to the Eurasian populations. For confirmation, we next applied the composite-likelihood-ratio (CLR) test within each population (Methods) and detected significant evidence of positive selection at the same locus in CHB and CEU, but not in YRI (Table S9), supporting the inference that positive selection acted in Eurasian populations likely after the out-of-Africa migration (see haplotype analysis for world-populations below).

Consistent with the XP-EHH results, the *NID2* promoter exhibited reduced haplotype diversity in both East Asian (CHB) and European (CEU) populations, as quantified by haplotype Shannon entropy (Methods; Fig. 4F, panel 2). This signature was substantially stronger in CHB, where the reduction in haplotype diversity coincided with a pronounced local minimum in recombination rate encompassing the locus (Fig. S5E), providing further evidence of positive selection. In CEU, both signatures were attenuated, potentially because demographic processes during the past 10,000 years partially eroded the earlier selection signal, as determined from ancient human genomes below. Collectively, these patterns are consistent with a selective sweep that increased the frequency of a beneficial allele or haplotype near the *NID2* promoter in Eurasia, with a more complete or better-preserved sweep in East Asians.

Using 1000 Genomes data, we identified 44 variants within this genomic region (Fig. 4F, and Table S10). Consistent with the East Asian-enriched XP-EHH signal, eleven of these variants were near fixation in East Asian populations, reached high population frequencies in Europeans and remained polymorphic in most African populations (Table S10). After assigning the evolutionarily ancestral and derived allele at each SNP (Methods), we evaluated the potential regulatory effect of each derived variant using our recently developed deep-learning framework.^46,47^ Specifically, using our single-nucleus ATAC-seq data as a DSC4 cell-state reference, the model achieved high predictive performance (Fig. S5F) and quantified the predicted allele-specific effects of each SNP within the *NID*2 promoter-flanking region on chromatin accessibility in DSC4 cells (Methods). This analysis identified SNP rs7154671, located upstream of the *NID2* transcription start site, as a candidate functional variant (Fig. 4F). The derived A allele was predicted to significantly reduce local chromatin accessibility relative to the ancestral G allele. Its frequency approached near fixation in East Asian populations and reached 89% and 51% in European and African populations, respectively (Fig. 4F, zoomed-in view; Table S10). We further examined rs7154671 allele frequencies across multiple independent population genomic resources, which consistently recapitulated the population-frequency pattern observed in the 1000 Genomes data (Fig. S6A).

To determine whether this predicted chromatin effect was associated with *NID2* expression in vivo, we first inferred rs7154671 genotypes for each individual from single-cell sequencing reads and then re-analyzed our single-nucleus RNA-seq data by stratifying individuals according to genotype and comparing NID2 expression in DSC4 cells across genotype groups (Methods). Subsequent eQTL analysis revealed a significant association between rs7154671 genotype and NID2 expression, with individuals homozygous for the derived A allele (A/A) exhibiting the lowest NID2 expression in DSC4 cells (Fig. 4G). This genotype-dependent expression pattern was consistent with the predicted effect of the derived A allele in reducing NID2 promoter accessibility (Fig. 4F). Together, these analyses identified rs7154671 as a candidate selected regulatory variant at the *NID2* promoter, in which positive selection fixed the derived allele in East Asians, resulting in reduced promoter accessibility and lower *NID2* expression in DSC4 cells.

*NID2* encodes nidogen-2, a secreted basement-membrane extracellular matrix protein. Given the role of DSC4 cells in suppressing EVT invasion,^4^ we reasoned that increased *NID2* expression by DSC4 cells may reinforce local basement-membrane and ECM organization at the decidual interface, thereby strengthening a stromal barrier that constrains EVT invasiveness. To test this possibility, we performed Matrigel transwell invasion assays using primary human cytotrophoblasts isolated from second-trimester samples. Cells were seeded onto Matrigel supplemented with either PBS control or recombinant NID2, thereby modeling increased extracellular NID2 deposition within the local matrix environment, and invaded cells were quantified after culture (Methods, Fig. 4H). Across three biologically independent experiments, each with four technical replicates, NID2 supplementation significantly reduced cytotrophoblast invasion compared with control (Fig. 4I), demonstrating that increased extracellular NID2 is sufficient to suppress trophoblast invasiveness. These findings support a model in which DSC4-derived NID2 contributes to an anti-invasive stromal matrix niche at the maternal–fetal interface (Fig. 4J). Accordingly, the reduced *NID2* expression associated with the East Asian-enriched derived allele may shift the decidual microenvironment toward greater permissiveness for EVT invasion, whereas higher *NID2* expression associated with the ancestral allele may confer stronger stromal restraint.

### Evolutionary origin of the NID2 promoter haplotype

Although both XP-EHH and CLR detected positive selection at the rs7154671 locus in European and East Asian populations, closer examination revealed distinct population-specific genetic architectures and regulatory consequences. First, DSC identity was negatively associated with East Asian ancestry, whereas the relative contribution of European versus African ancestry (conditional on East Asian ancestry) was not significantly associated with DSC identity (Table S8). Second, the local selection signature was substantially stronger in East Asians: European populations retained greater haplotype diversity and higher local recombination rates than East Asian populations (Fig. 4F; Fig. S5E), indicating a less complete or more extensively eroded selective sweep. Third, and most notably, we identified a nearby variant, rs12886723, whose derived allele was predicted to significantly increase local chromatin accessibility, opposing the accessibility-reducing effect of the derived allele at rs7154671 under positive selection (Fig. 4F). This rs12886723-derived allele was strongly enriched in European populations, occurring at a frequency of approximately 10% but remaining rare in other populations (Fig. 4F).

To distinguish these opposing regulatory effects, we designate the evolutionarily selected rs7154671 as locus A: its ancestral and derived alleles are A^Anc+^ and A^Der−^, respectively, reflecting increased and decreased local chromatin accessibility. Accordingly, we designate rs12886723 as locus B: its ancestral and derived alleles are B^Anc−^ and B^Der+^ for their respectively predicted allelic effects on chromatin openness. In human populations, we found that the accessibility-increasing B^Der+^ allele was consistently linked to A^Anc+^, forming an A^Anc+^B^Der+^ haplotype that functionally opposes the selected, accessibility-reducing A^Der−^B^Anc−^ haplotype. European populations therefore contain two major regulatory haplotypes: A^Der−^B^Anc−^ predicted to reduce *NID2* expression, and the Europe-specific A^Anc+^B^Der+^, predicted to increase NID2 expression (accounting for 10.5% of the population). Their worldwide distributions are illustrated by the allele frequencies of rs7154671 (locus A) and rs12886723 (locus B) (Fig. 5A).

**Figure 5.**
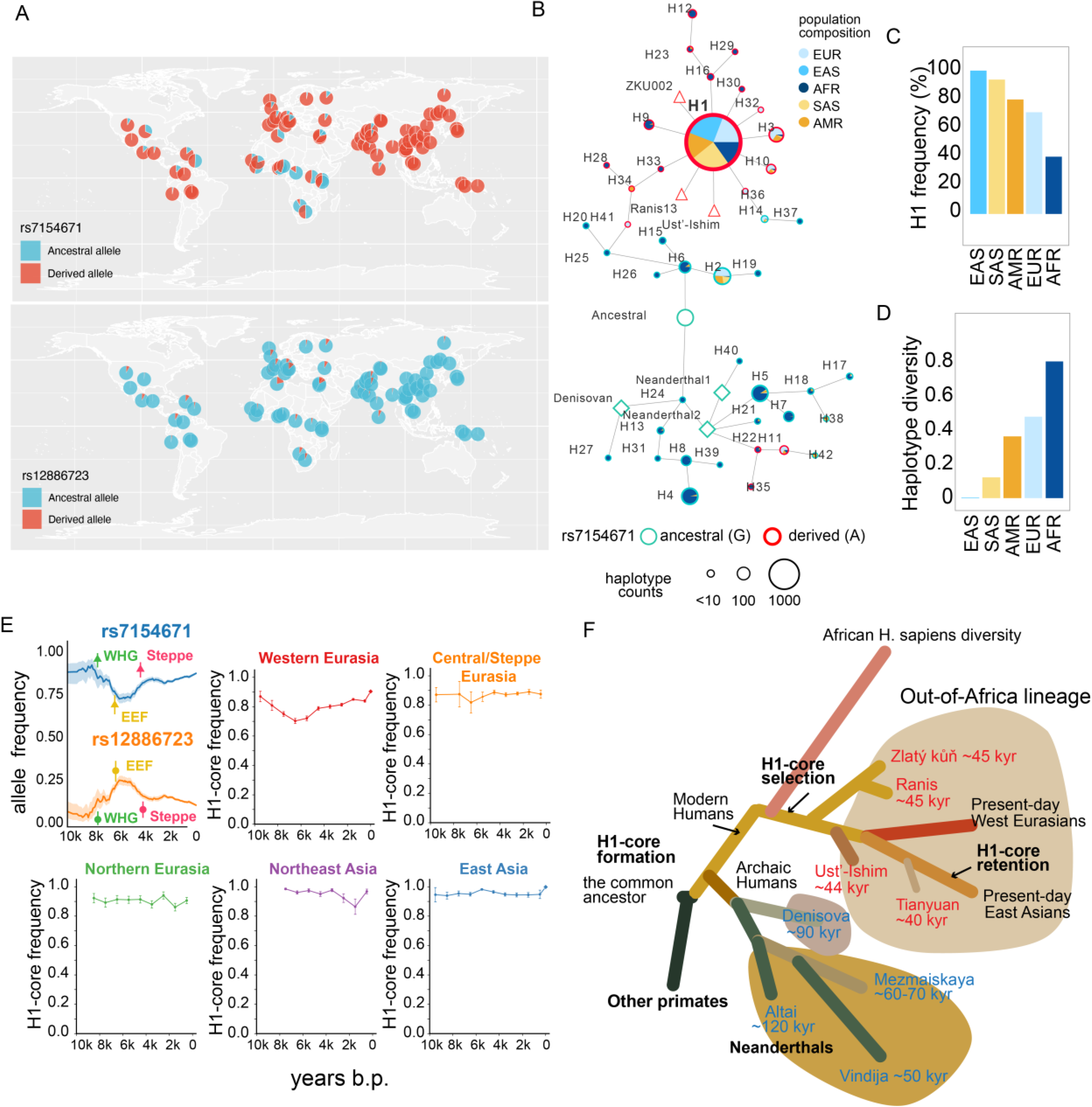
Global distribution, haplotype structure, and evolutionary history of the NID2 associated haplotypes. (A) Geographic distribution of ancestral and derived alleles at rs7154671 and rs12886723 across worldwide populations. (B) Haplotype network of the NID2 promoter region. Nodes represent distinct haplotypes, with node size indicating haplotype frequency in the overall modern human population and node color indicating the geographic superpopulation composition defined by the 1000 Genomes Project. Edges represent mutational differences between haplotypes. The ancestral and derived allelic states at rs7154671 are indicated by green and red node outlines, respectively, and selected ancient modern human and archaic genomes are labeled. (C) Frequency of the H1 haplotype across continental ancestry groups. (D) Haplotype diversity across the same populations. (E) Temporal patterns of the focal variants and H1-core two-locus configuration in ancient Eurasian populations. The first panel shows AGES estimates for rs7154671 and rs12886723 separately, with WHG (Western hunter-gatherers), EEF (Early European Farmers), and Steppe-related groups indicated. The remaining panels show reconstructed temporal patterns of the H1-associated rs7154671-A/rs12886723-A two-locus configuration across five Eurasian regions using AADR ancient individuals. (F) Schematic reconstruction of the evolutionary history of the H1-core two-locus configuration, illustrating its formation in modern humans, subsequent increase in frequency after the out-of-Africa expansion, retention in present-day West and East Eurasian populations, and relationships to representative archaic and ancient human lineages.

To reconstruct the evolutionary history of this architecture, we analyzed promoter-proximal haplotypes centered on rs7154671 (locus A) in 3,202 individuals with phased genomes from the 1000 Genomes Project Phase 3 panel. We identified 42 distinct haplotypes (Table S11; Methods) and reconstructed their relationships in a haplotype network (Fig. 5B), and examined the global distribution of major haplotypes across populations (Fig. S6B). H1, the predominant *NID2*-reducing haplotype carrying A^Der−^B^Anc−^ at the haplotype core, was nearly fixed in East Asians, compared with frequencies of 70.9% in Europeans and 39.9% in Africans (Fig. 5C). As expected, this population-frequency gradient was accompanied by a stepwise reduction in local haplotype diversity, from 0.79 in Africans and 0.47 in Europeans to approximately 0 in East Asians (Fig. 5D). The A^Der−^B^Anc−^ two-locus configuration was also present in approximately 40,000–45,000-year-old modern Eurasians, including Zlatý kůň, Ranis13, and Ust′-Ishim (Fig. S6C-D). In the haplotype network, all haplotypes carrying the *NID2*-reducing A^Der−^ allele formed a clade distinct from haplotypes related to Neanderthals and Denisovans (Fig. 5B), indicating that this regulatory allele arose specifically on the modern-human lineage. Also notably, the A^Der−^ allele appeared before formation of the complete H1 haplotype, as indicated by additional A^Der−^-carrying haplotypes (red-bordered nodes in Fig. 5B).

To date the focal mutations, we reconstructed local genealogies with Relate^48^ using 310 present-day individuals (99 CEU, 103 CHB, and 108 YRI) and four high-coverage archaic genomes: the Altai, Vindija 33.19, and Chagyrskaya 8 Neanderthals and the Denisova 3 Denisovan. The A^Der−^ mutation at rs7154671 (locus A) mapped to a modern-human branch dated to approximately 306–383 thousand years ago (Methods), consistent with its absence from the archaic genomes and its presence in early modern Eurasians (Zlatý kůň, Ranis13, and Ust′-Ishim, Fig. 5B; Fig. S6D). Therefore, the shared Eurasian selection signature most likely reflects positive selection acting on pre-existing standing variation after the out-of-Africa migration but before the divergence of West and East Eurasian populations.

By sharp contrast, the accessibility-increasing B^Der+^ allele at rs12886723 (locus B, Fig. 4F) occurred together with the ancestral allele at rs7154671, forming the A^Anc+^B^Der+^ two-locus configuration. This configuration was distributed across several haplotypes but was predominantly carried by H2 (Fig. 5B, Table S11). This principal mutation-bearing branch was dated to 14.6–27.8 thousand years ago (Methods), suggesting very recent origin and subsequent expansion only in European populations. We detected no clear positive-selection signature around this locus (locus B), suggesting that its present-day European enrichment may instead reflect recent demographic history or genetic drift, which may also have weakened the older selection signal around rs7154671 (locus A) as discussed above.

Consistent with this, we queried the AGES database,^49–51^ which recorded the estimated allele frequencies in Western Eurasia over the past 10,000 years through ancient human genome, and observed that the frequencies of these two antagonistic alleles (A^Der^- and B^Der+^) changed markedly over the past 10,000 years (Fig. 5E). The selected A^Der^-allele declined from approximately 90% to 70–75% during the early-to-middle Holocene before recovering to approximately 85% in present-day Europeans. The opposing B^Der+^ allele rose from approximately 5% to 25% before declining to its present-day frequency of approximately 10–15%. These shifts closely tracked changes in European ancestry composition: A^Der−^ was common and B^Der+^ rare in Western hunter-gatherers (WHG) and Steppe-associated populations, whereas Early European Farmers (EEF) showed the opposite pattern (Fig. 5E). Thus, the present-day frequencies of these antagonistic regulatory alleles likely reflect repeated demographic turnover and ancestry admixture superimposed on, and partially obscuring, the effects of selection prior to West and East Eurasian population split.

To determine whether this instability extended across Eurasia, we analyzed 7,942 ancient individuals from the Allen Ancient DNA Resource (AADR)^52^, including 319 from East Asia, 279 from Northeast Asia, 529 from Northern Eurasia, 1,239 from Central Eurasia or the Steppe, and 5,576 from Western Eurasia. Because many ancient genomes lacked direct coverage of the two focal SNPs (locus A and B), we inferred the probability of carrying the H1 core two-locus configuration (A^Der−^B^Anc−^) from 80 harmonized biallelic flanking markers spanning the union of the 100-kb windows surrounding both loci (Methods). The inference model closely reproduced directly observed haplotype frequencies in modern populations (Table S12) and was then applied to the AADR genomes. Consistent with individual SNPs (Fig. 5E), Western Eurasia showed the greatest temporal fluctuation in the two-locus *NID2* regulatory configuration (Fig. 5E), which, in contrast, remained near fixation in East Asia throughout the past 10,000 years (Fig. 5E). We further applied this strategy to the Tianyuan individual, an ancient East Asian dated to approximately 40,000 years ago,^53^ and predicted with high confidence that the Tianyuan individual was homozygous for the H1-associated two-locus configuration (Table S12). Thus, NID2-reducing haplotype likely has been maintained at a very high frequency (near fixation) in East Asian-related populations for at least 40,000 years. Meanwhile, Northern Eurasia and Northeast Asia showed intermediate patterns, thereby defining a broad west-to-east cline of progressively greater retention of the NID2-reducing haplotype toward East Asia (Fig. 5E).

Taken together, these population trajectories connect the evolutionary history of the *NID2* promoter-proximal regulatory region to its functional effects on reducing EVT invasion. In East Asian-related populations, H1 remained near fixation throughout the past 10,000 years, consistent with sustained retention of the selected *NID2*-reducing haplotype. Because DSC4-derived *NID2* restrains fetal EVT invasion, H1 is predicted to reduce stromal restraint and create a uterine microenvironment more permissive to EVT invasion; its near fixation in East Asia therefore suggests that selection may have strongly favored reduced NID2-mediated restraint and thus a more EVT-permissive uterine microenvironment. In Western Eurasia, repeated population turnover, together with the emergence of the *NID2*-increasing A^Anc+^B^Der+^ allelic configuration, generated substantially greater heterogeneity in predicted *NID2* expression. This heterogeneity likely weakened the correspondence between European ancestry and both DSC identity and NID2 expression, thereby obscuring the present-day regulatory consequences of past positive selection even though its genomic signature remains detectable. In African populations, by contrast, the H1 haplotype occurred at a substantially lower frequency (39.9%), and no significant positive-selection signal was detected at this locus in YRI, consistent with retention of greater ancestral haplotypic diversity. This broader genetic variation is expected to generate greater interindividual variation in *NID2* regulation and may therefore contribute to the greater heterogeneity of pregnancy-related phenotypes observed across individuals of African ancestry. These population-specific trajectories of this modern-human-specific regulatory innovation are briefly summarized in Fig. 5F.

## Discussion

These single-cell analyses showed that human placentation uses largely the same trophoblast differentiation programs across the entire maternal-fetal interface, which unfolds differently in the CM and BP, resulting in their distinct tissue architectures. CM builds a laminar trophoblast barrier in which EVT invasion is constrained. In contrast, EVTs in the BP have evolved into subpopulations that carry out a variant of collective migration, which is observed in tumor metastasis. Invasion is further tuned by a maternal stromal rheostat, *NID2* in DSC4 cells, whose activity varies across human populations and appears to have been modified by recent positive selection. Together, the results of this study reveal human placentation as an evolutionarily tunable system in which overlapping differentiation programs give rise to conserved trophoblast states that are deployed differently in the BP vs. the CM with additional input from population-variable decidual control of EVT invasion.

Our findings suggest modification of the conventional linear model in which EVT maturation is assumed to coincide with progressively deeper invasion. Instead, the location and phenotypes of early-arising and mature EVTs suggest a nonlinear model. Early-arising EVTs mediate deep decidual and arterial invasion, whereas mature EVTs remodel the superficial immune and stromal interface. It is possible that the deep decidual microenvironment contributes to the long-term maintenance of the immature status of EVTs for long-range migration at the invasive front. As such, these deep EVTs likely culminate in invasion of the muscular layer of the uterus, a theory we could not investigate as our samples did not include placental bed biopsies. Regardless, this framework suggests that pregnancy disorders associated with abnormal invasion could arise from either of the identified EVT subpopulations, a combination of both or, more important, an imbalance between these coordinated EVT states.

Human EVTs in the BP are often viewed through the lens of immune evasion, with the nonclassical MHC class I molecule, HLA-G, enabling embryonic/fetal cell invasion of the uterus while avoiding immune rejection. Our data showed that the canonical EVT immune program— including the expression of *HLA-G*, *HLA-C*, *B2M*, and *TRAIL*—was upregulated in mature EVTs enriched in the superficial decidua. Thus, mature EVTs may couple MHC class I-mediated immune tolerance with TRAIL-dependent elimination of select decidual immune populations, shaping the local immune microenvironment to create a permissive niche for continued trophoblast expansion and occupancy. In contrast, the early-arising EVTs that penetrate the deep decidua express markedly reduced levels of these immune-modulatory molecules, including *HLA-G* (Fig. 3M). Given the dense immune-cell enrichment in the deep decidua,^4^ this finding suggests that deep EVT invasion is unlikely to be facilitated by the same mechanisms employed in the superficial decidua. Instead, tolerance of the early arising EVTs in the deep decidua may involve a specialized immune-compatibility program, a mechanism that warrants further investigation.

EVT penetration of the uterus is often compared with tumor invasion, but this analogy has lacked more specific details. Based on the data presented here, we theorize that the early-arising and deeply invading EVTs have functions akin to the leader cells that initiate tumor cell metastasis, one example of collective cell migration. In the decidua, these EVTs activate a navigation-centered program with striking parallels to neuronal migration, including the extension of polarized protrusions and the use of canonical guidance pathways such as ephrin^54^ and semaphorin signaling.^55^ Our in vitro imaging previously showed that invasive EVTs extend long processes ahead of the cell body,^56^ consistent with exploratory pathfinding behavior that could interpret spatial guidance cues that direct them to arterial targets and the deep decidua. Meanwhile, mature EVTs appear to act as local niche engineers, remodeling the extracellular matrix and sculpting the immune microenvironment of the maternal-fetal interface to create permissive routes for deep invasion by the early-arising EVTs.

The data presented here also enabled new insights into formation of the CM. We found evidence that ghost villi, regressed chorionic villi that were once in touch with the uterine wall, may have entered an anchoring villus developmental program before involution. Consistent with this theory, the basal layer of the trophoblast shell was formed by a uniform layer of EVT progenitor-like cells. In the BP, these cells are restricted to cytotrophoblast cell columns of anchoring villi. Moreover, the fetal mesenchymal compartment underlying the EVT shell contained Hofbauer-like fetal macrophages and fibroblasts, which are key villous components. Together, these observations suggest that, following regression of membrane-associated villi, their mesenchymal cores and associated cell columns may laterally expand into the planar architecture of the CM.

The dominance of DSC4 cells adjacent to the CM was unexpected. We had initially identified this population in the BP, where they are sparsely distributed across the superficial decidua and likely restrain invasive EVTs that emerge from cytotrophoblast cell columns.^4^ In sharp contrast, DSC4 cells in the CM formed a dense layer apposed to the cytotrophoblast compartment. We theorize that they restrict EVT migration thereby helping to retain the epithelial-like integrity of the CM.

Our study also places the DSC4-NID2 axis within the evolutionary history of human placentation. Across primates, deep interstitial cytotrophoblast invasion was newly established in the great-ape lineage.^57^ Thus, the selection of the NID2 promoter haplotype in modern humans suggests a very recent innovation that further modifies placental invasion by increasing the permissiveness of the uterine microenvironment. Furthermore, this strategy was absent from Neanderthal and Denisovan archaic humans but was strongly favored by East Asian populations. Such tuning could have enhanced reproductive fitness by supporting implantation, EVT invasion, and consequently, fetal growth, which would ward off complications such as severe preeclampsia, growth restriction and some cases of preterm birth. The tradeoff may be increased vulnerability to excessive or mislocalized invasion, including placenta accreta spectrum (PAS). Indeed, epidemiological data show lower rates of preeclampsia^58,59^ and preterm birth^60,61^ but higher rates of PAS^62^ (even in the absence of prior cesarean deliveries^63^) among East Asian populations.

Although excessive trophoblast invasion can be pathological, insufficient invasion likely imposed a more immediate and recurrent constraint on reproductive fitness by compromising placental anchoring, vascular remodeling, fetal growth, and pregnancy maintenance. This tradeoff may have been especially consequential during the eastward dispersal of modern humans, when ancestral East Asian populations experienced serial founder effects, stronger demographic bottlenecks, and smaller effective population sizes. Under such conditions, even modest gains in pregnancy maintenance and offspring survival could generate substantial fitness advantages. Thus, selection may have favored a *NID2* promoter haplotype that shifted the uterine microenvironment toward greater permissiveness for fetal invasion, because the net reproductive gain from reducing insufficient placentation outweighed the later or less frequent evolutionary cost of excessive invasion.

However, although our analyses emphasize population-level ancestry patterns, the functional consequences of this variation ultimately operate at the level of individual genomes. Even in populations without evidence of positive selection at this locus, the *NID2*-reducing A^Der−^ allele remains widespread: for example, occurring at a frequency of 39.9% in African populations. Its broad and polymorphic distribution is therefore expected to generate substantial interindividual variation in *NID2* expression, which may contribute, at least in part, to the marked heterogeneity in susceptibility to pregnancy complications both within and across populations.

### Study Limitations

Several limitations remain. First, our cohort was restricted to clinically uncomplicated pregnancies, which allowed us to define population-associated baseline variation at the maternal-fetal interface, but does not directly establish how these baseline differences translate into disease. Future studies will need to examine clinical cohorts, including our companion severe preeclampsia and preterm birth study, to determine whether ancestry-associated shifts in DSC4, NID2 regulation, and EVT invasion programs are amplified or disrupted in pregnancy complications associated with shallow placentation. Second, many ancestry-dependent effects on placentation are likely to manifest during early gestation, initially when implantation and trophoblast lineage allocation occur, and later, when EVT invasion and spiral artery remodeling take place in a permissive decidual environment. Because these seminal stages of early pregnancy were not captured in the present study, future profiling of these events will be essential to determine when population-variable uterine microenvironments first emerge and how they shape later pregnancy outcomes.

## Acknowledgements

This work was supported by the Chan-Zuckerberg Initiative (the Silicon Valley Community Foundation, DAF2021-239932 and DAF2020-217686) and the National Institute of General Medical Sciences (R35GM142983) to J.L. X.S. discloses support for the research of this work from the National Institute of Child Health and Human Development (NICHD) (R01HD068524). D.K.S. was supported by Christopher Hess Research Fund (Stanford University). S.K.E. was supported by the March of Dimes Fund (Prematurity Research Center, Washington University). This work was supported in part by The Reproductive Specimen Processing and Banking Biorepository at Washington University School of Medicine.

