## Supplementary figures and images for "Spatial Logic and Evolutionary Innovation in Human Placentation"

### Supplemental Figure 1

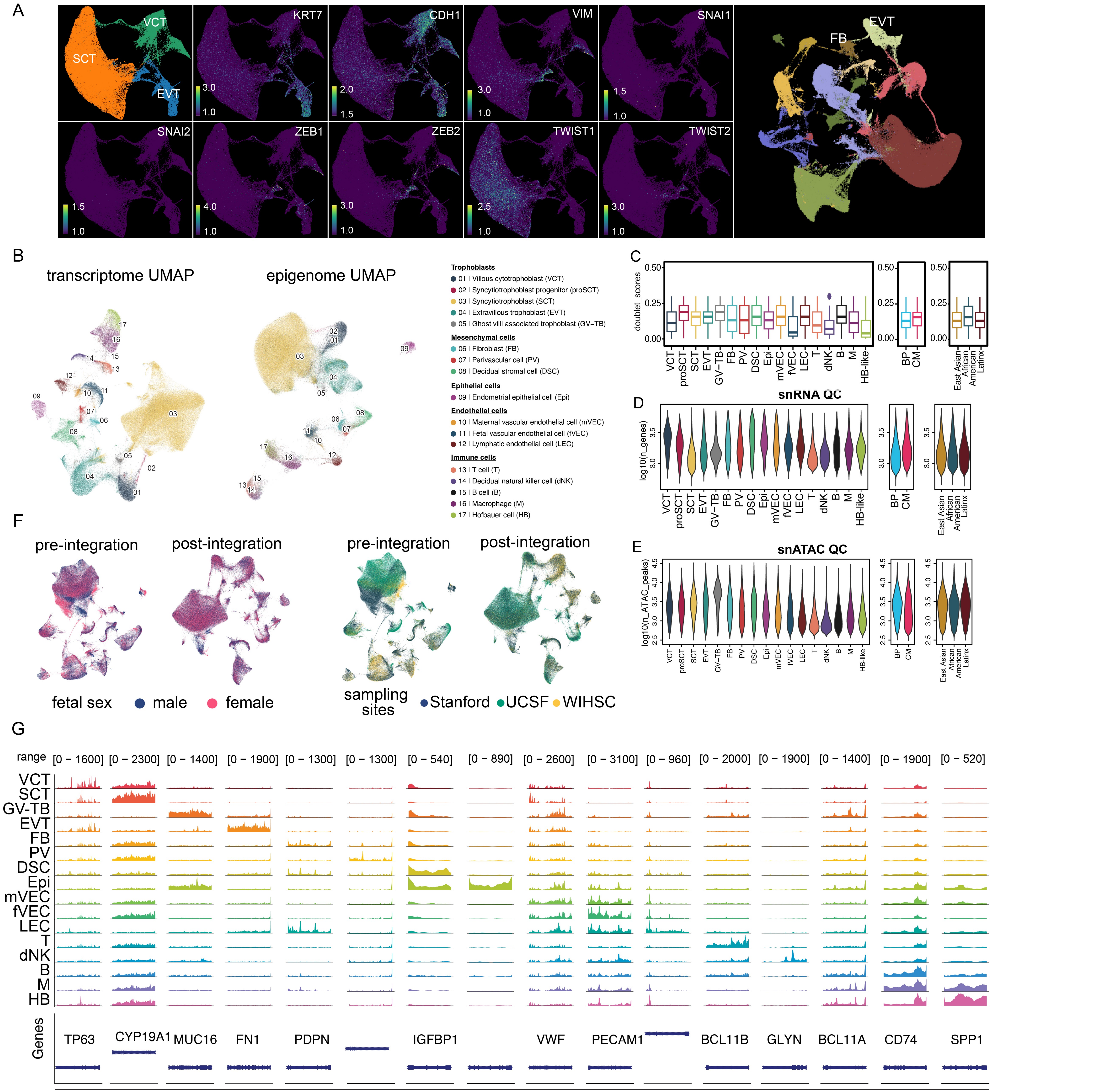

### Supplemental Figure 2

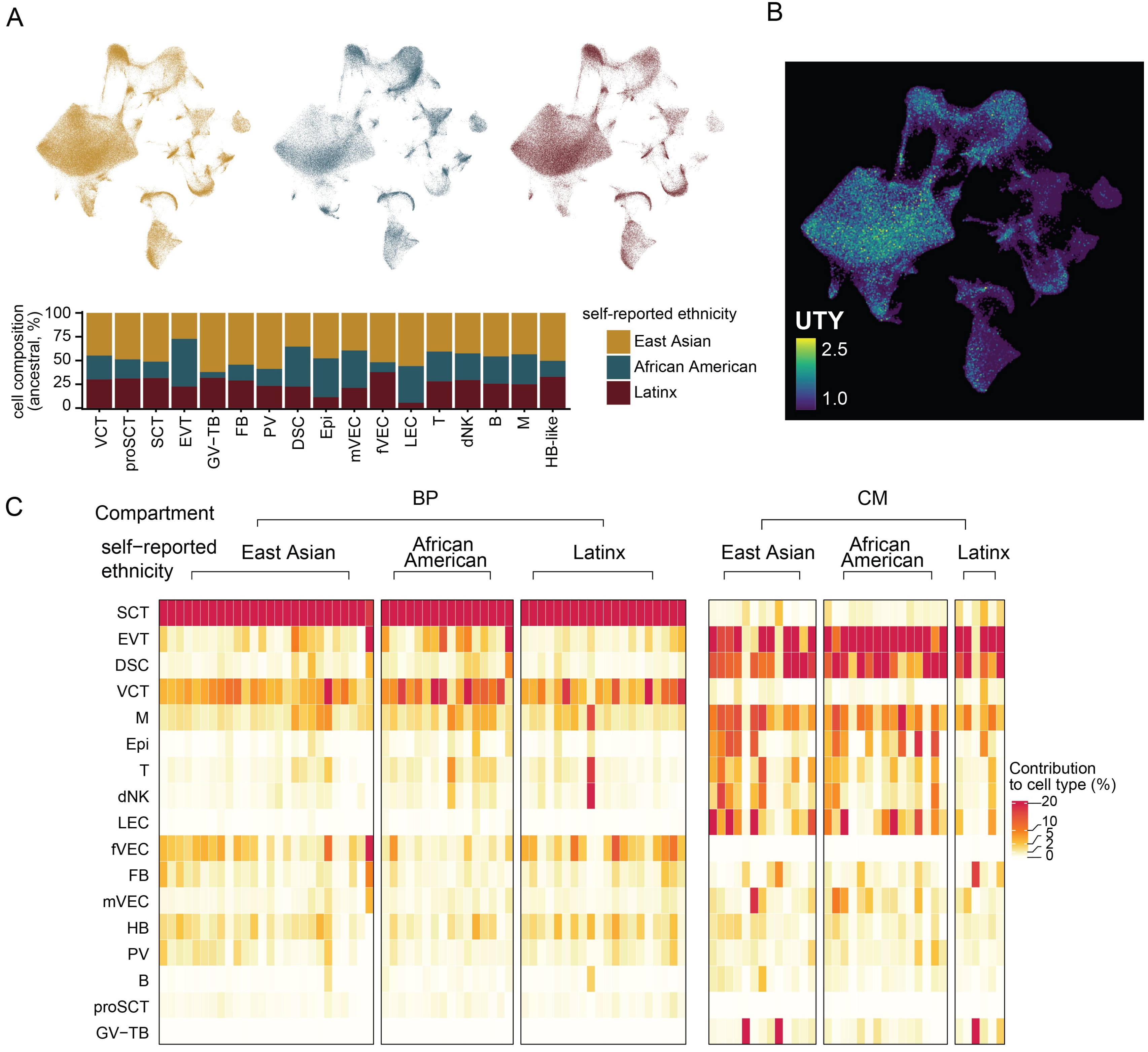

### Supplemental Figure 3

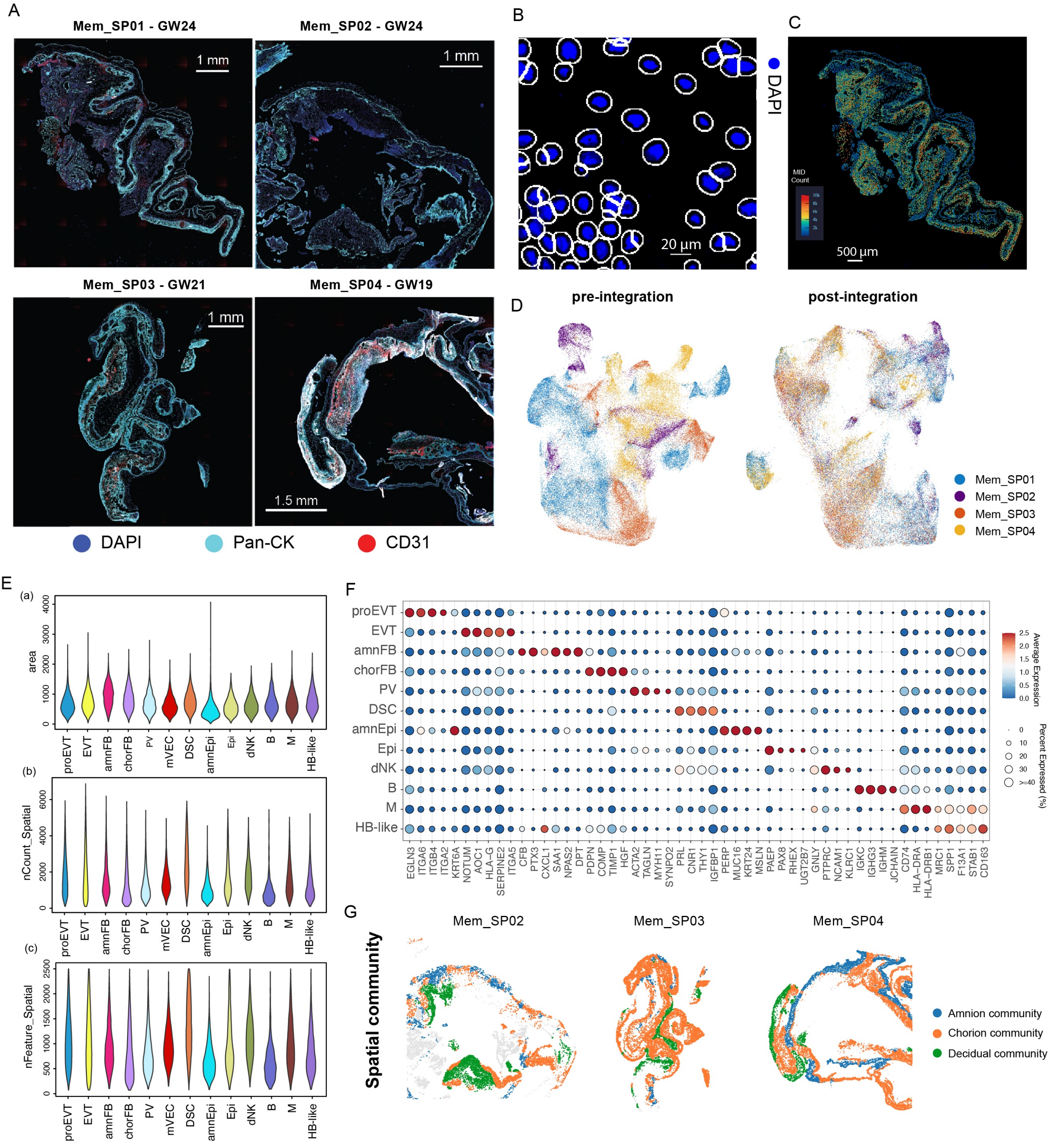

### Supplemental Figure 4

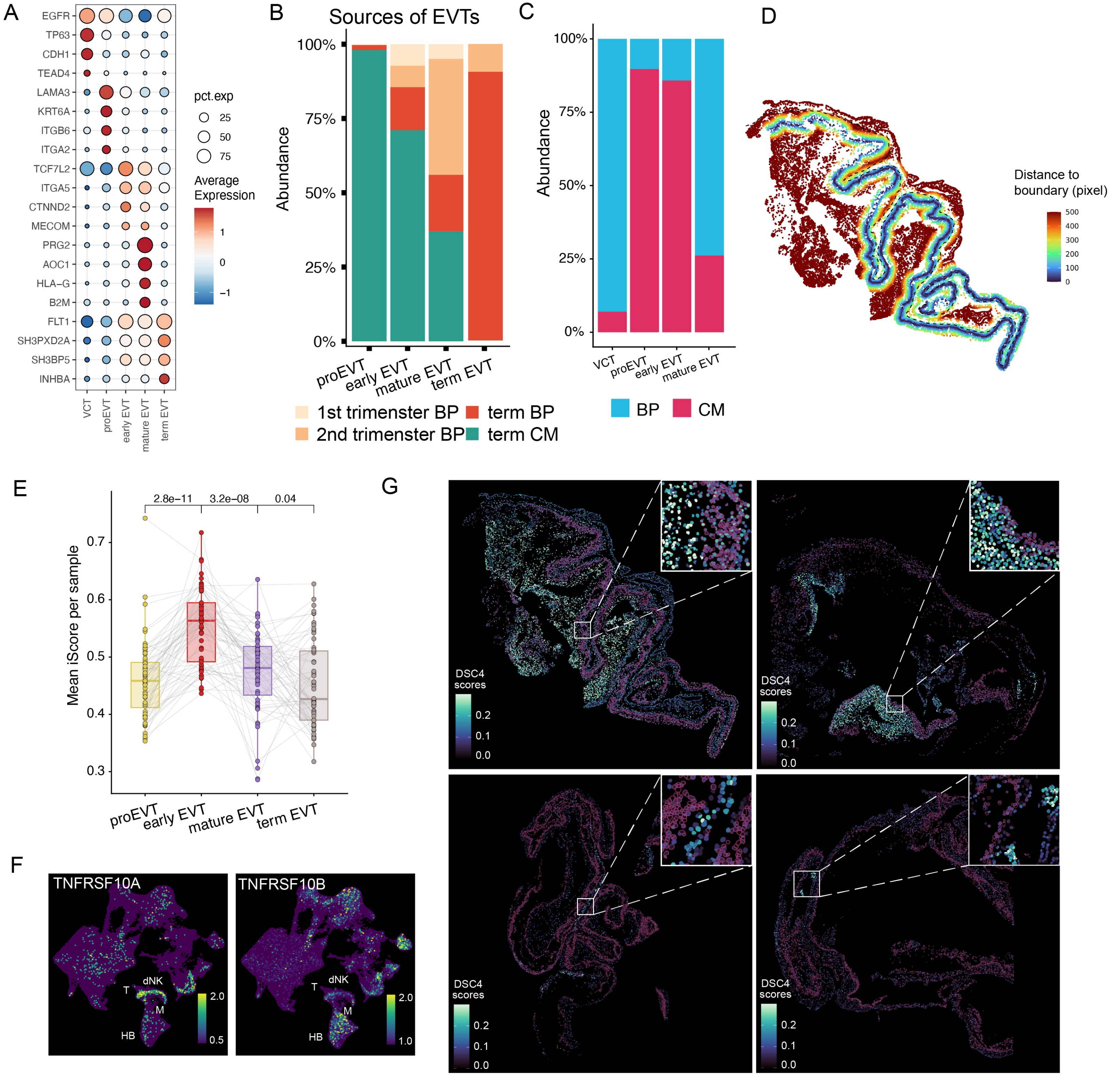

### Supplemental Figure 5

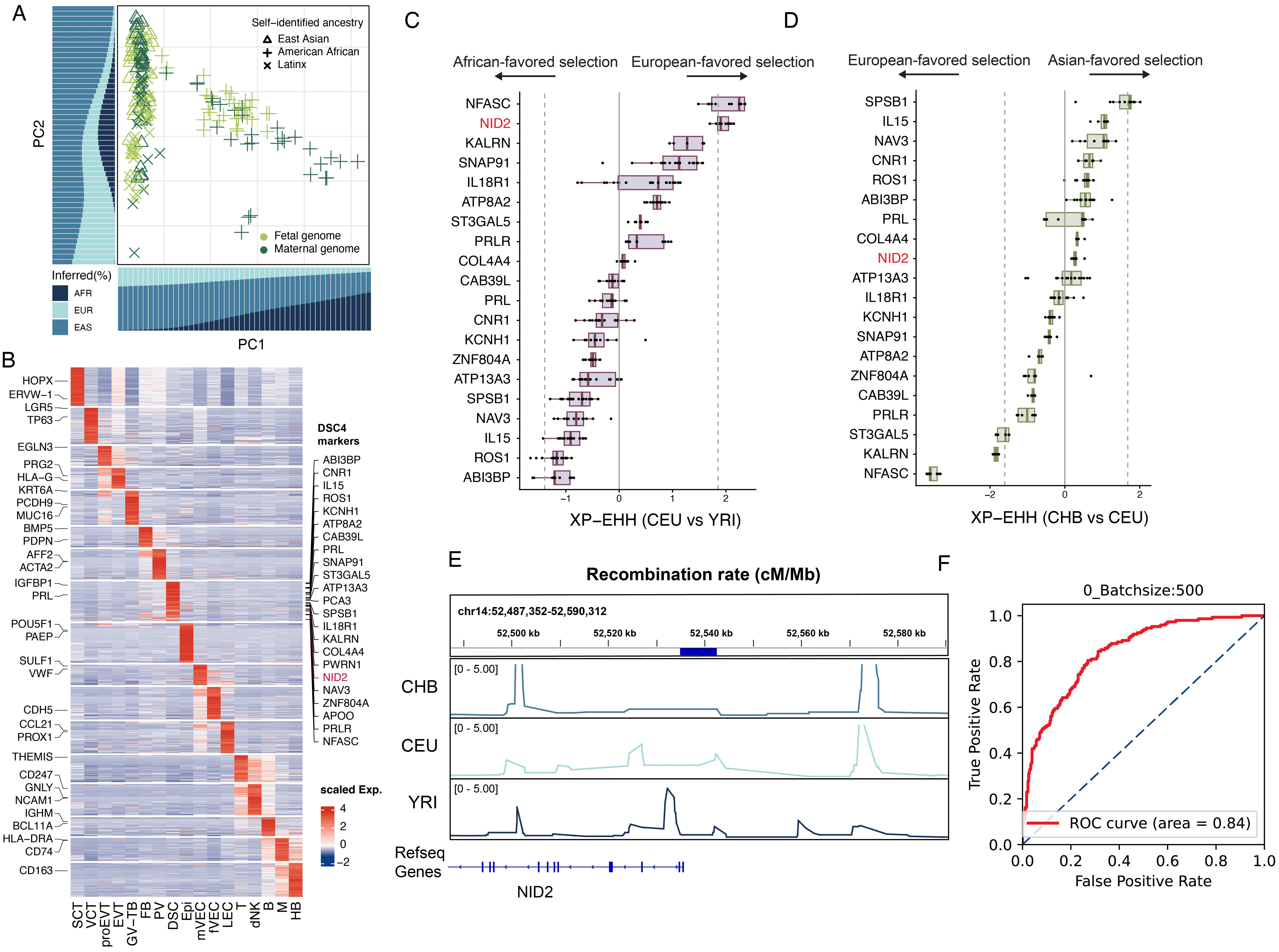
