## Supplemental Figure 6 for "Spatial Logic and Evolutionary Innovation in Human Placentation"

A

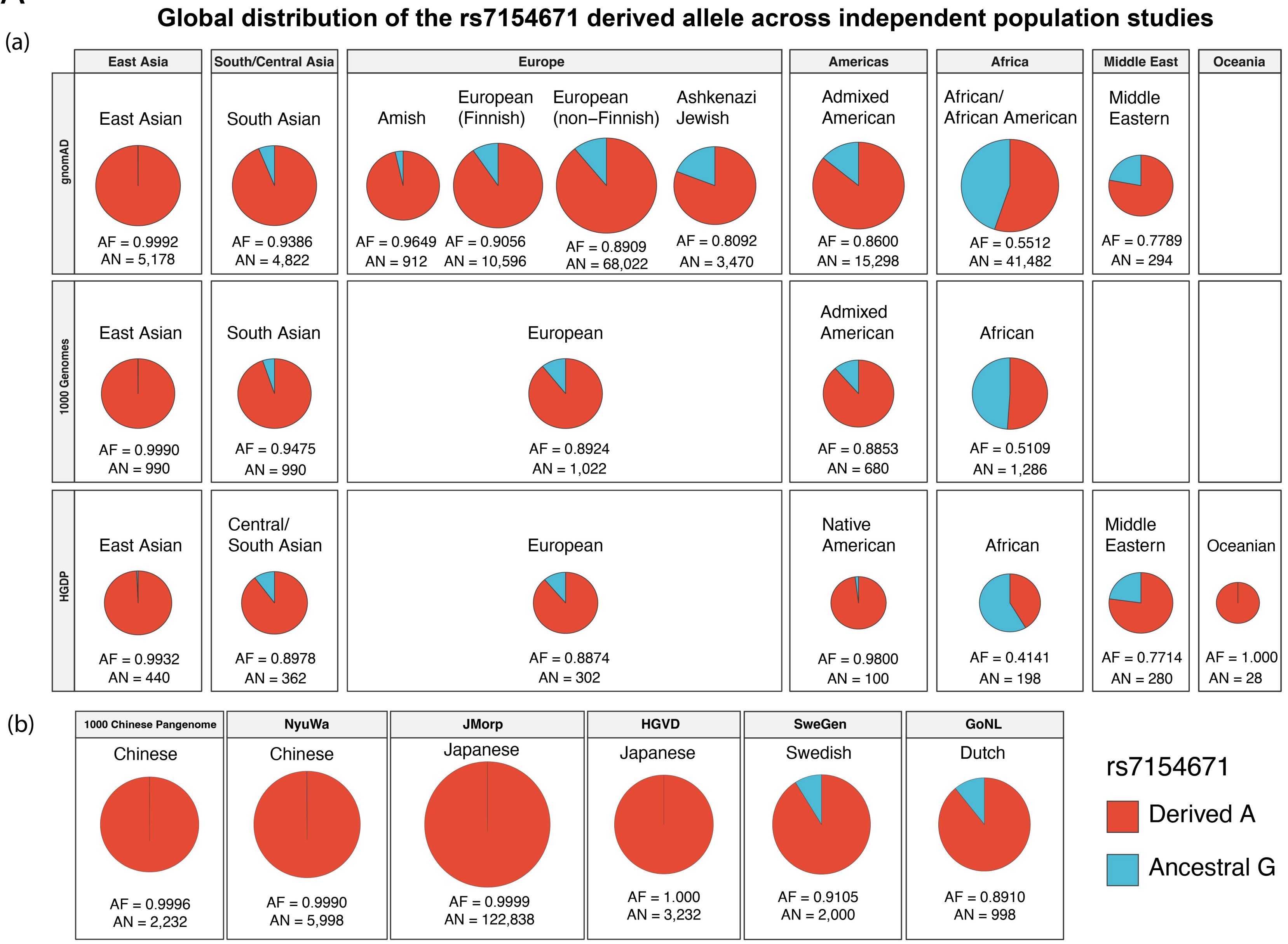


B

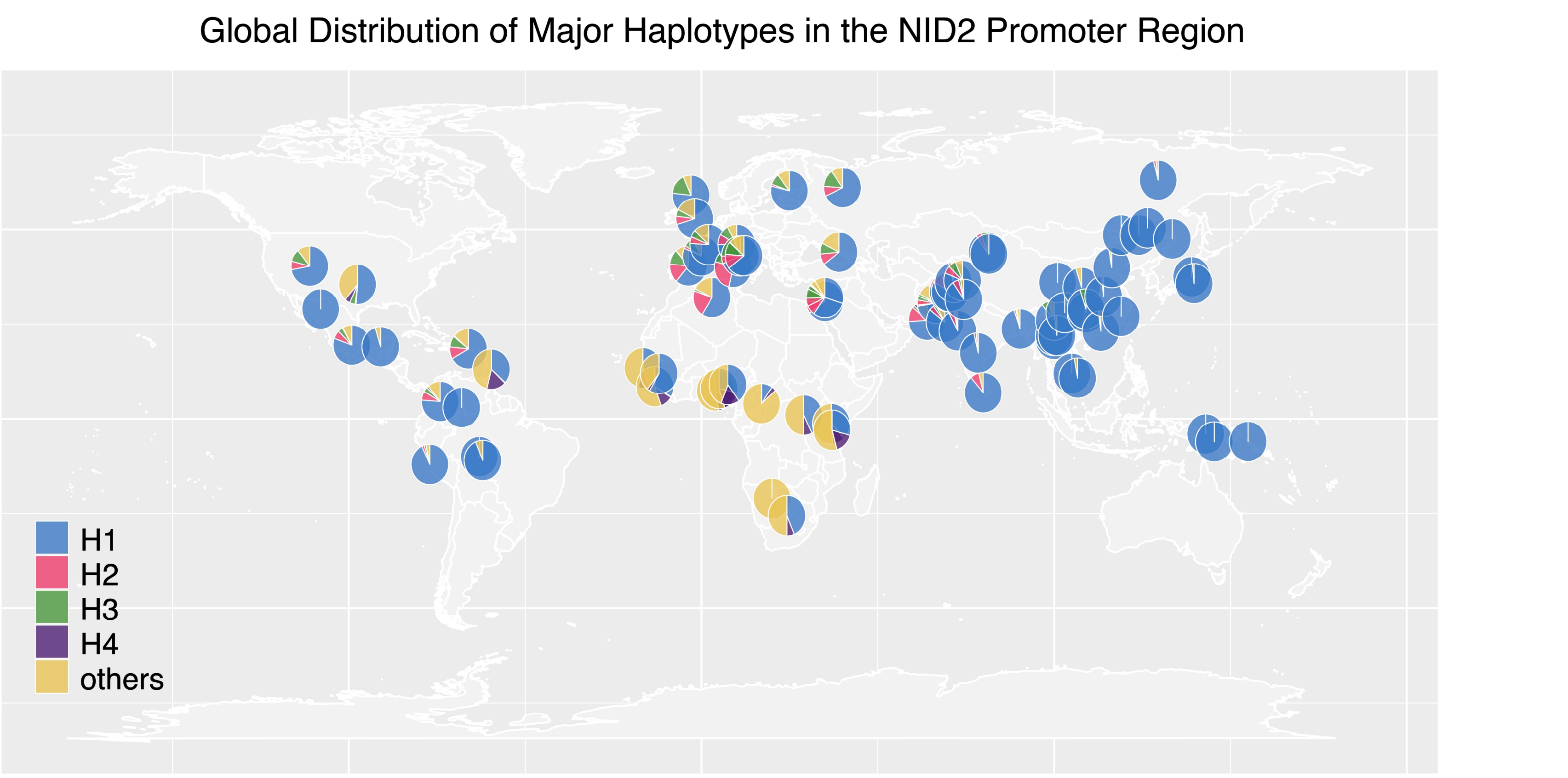


C

| Sample | Lineage | Age (kya) | Region | Coverage |
|---|---|---|---|---|
| Ust'-Ishim | Early modern human | ~45 | Western Siberia, Russia | High |
| Ranis13 | Early modern human | ~45 | Ranis, Germany | High |
| ZKU002 | Early modern human | >45 | Zlatý kůň, Czechia | High |
| Mezmaiskaya 1 | Neanderthal | ~60–70 | Northern Caucasus, Russia | Low |
| Altai Neanderthal | Neanderthal | ~120 | Denisova Cave, Altai, Russia | High |
| Chagyrskaya 8 | Neanderthal | ~80 | Chagyrskaya Cave, Altai, Russia | High |
| Vindija 33.19 | Neanderthal | ~50–52 | Vindija Cave, Croatia | High |
| Denisova 3 | Denisovan | ~65–72 | Denisova Cave, Altai, Russia | High |

D

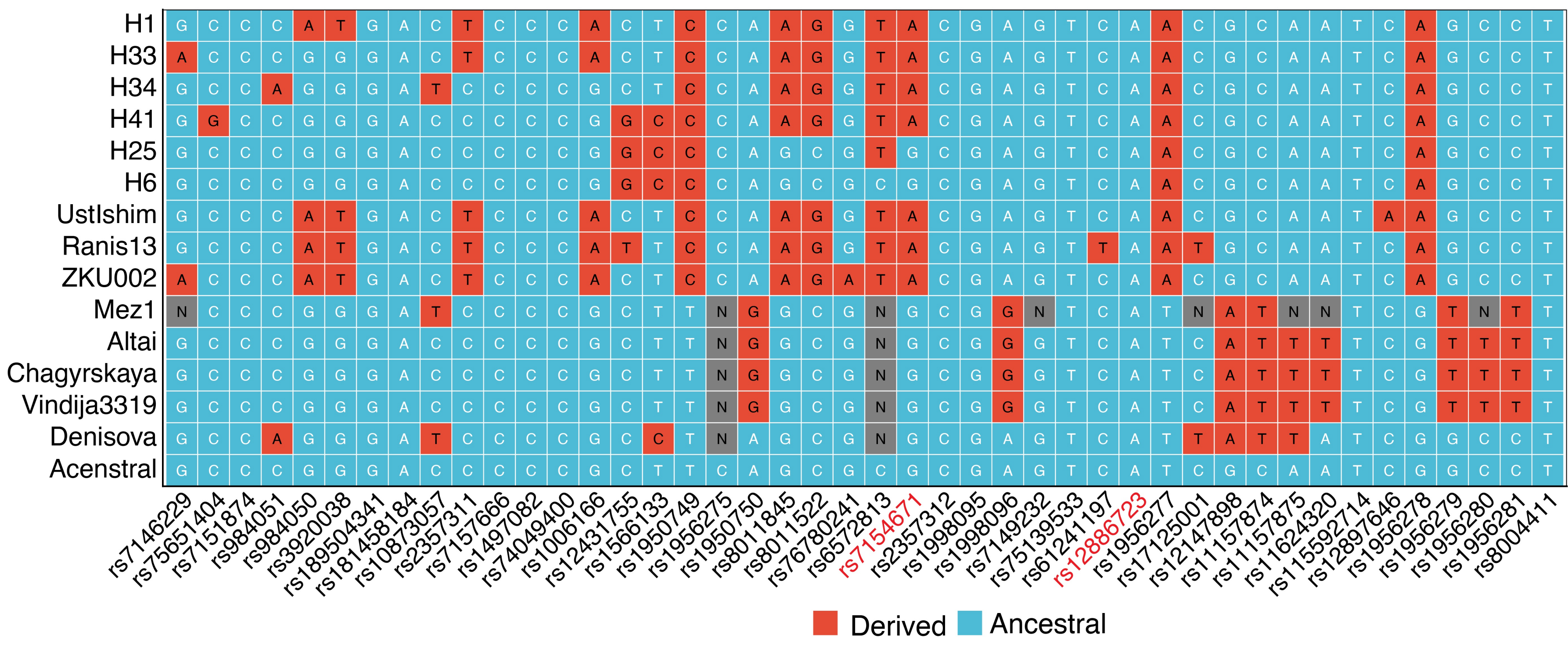
